# Release the Krakencoder: A unified brain connectome translation and fusion tool

**DOI:** 10.1101/2024.04.12.589274

**Authors:** Keith W. Jamison, Zijin Gu, Qinxin Wang, Ceren Tozlu, Mert R. Sabuncu, Amy Kuceyeski

## Abstract

Brain connectivity can be estimated in many ways, depending on modality and processing strategy. Here we present the Krakencoder, a joint connectome mapping tool that simultaneously, bidirectionally translates between structural (SC) and functional connectivity (FC), and across different atlases and processing choices via a common latent representation. These mappings demonstrate unprecedented accuracy and individual-level identifiability; the mapping between SC and FC has identifiability 42-54% higher than existing models. The Krakencoder combines all connectome flavors via a shared low-dimensional latent space. This “fusion” representation i) better reflects familial relatedness, ii) preserves age- and sex-relevant information and iii) enhances cognition-relevant information. The Krakencoder can be applied without retraining to new, out-of-age-distribution data while still preserving inter-individual differences in the connectome predictions and familial relationships in the latent representations. The Krakencoder is a significant leap forward in capturing the relationship between multi-modal brain connectomes in an individualized, behaviorally- and demographically-relevant way.

## 1 Introduction

The brain is a vastly complex, interconnected network of neurons (and other types of cells) whose proper function allows us to move, think, feel, and observe as well as interact with our environment. Studying how the brain’s connections relate to behavior, so-called brain connectome-behavior mapping, is crucial not only for understanding how the brain works generally but also for identifying biomarkers of disease, predicting outcomes in neurological disorders, and designing personalized interventions. Brain connectivity can be probed through various structural and functional neuroimaging techniques. Structural connectivity (SC) can be captured with diffusion MRI (dMRI) measures of white matter, or the anatomical wiring, that connects different brain regions. Functional connectivity (FC), as measured with blood oxygen level dependent functional MRI (BOLD-fMRI), quantifies the temporal similarity between brain regions’ dynamic neural activity patterns.The association between SC and FC is a subject of intense research, as it is believed that structural pathways provide the backbone over which functional activation flows. Despite this, correlations of SC and FC are moderate at best. Understanding the relationship between SC and FC is key to deciphering how the brain’s physical structure supports its dynamic functions and may offer insights into the neural mechanisms underlying behavior, injury or disease, and recovery^1–4^.

There are many approaches to modeling the relationship between SC and FC, most of which focus on using SC to predict FC. These methods include i) biophysical models, e.g. linked neural mass models^5–10^, ii) graph theoretical models, e.g. communication strategies^11–15^, iii) statistical models, i.e. that model SC and FC topological metrics^16–19^, and iv) network control theory^20–23^. More recently, machine learning has been used to predict FC from SC^24–27^, predict SC from FC^28^, or to translate a connectivity estimate between parcellations^29, 30^. Many of these studies have shown that metrics of SC-FC relationships vary with age, sex, and behavioral variables^16, 27, 31^, or disease/injury and recovery^4, 32, 33^.

Among the challenges and considerations in modeling this relationship, the assumption that estimated SC is an objective, fixed constraint on possible FC patterns is increasingly questioned. Evidence suggests that the brain’s functional dynamics can be highly flexible and adaptive, and may be driven by geometric factors not measured with SC^34^. Variability in SC estimation techniques, particularly in their sensitivity to detect faint or indirect connections, further complicates the interpretation of connectivity data. Moreover, the performance metrics and loss functions currently employed in connectivity studies often fail to adequately account for inter-subject variability, potentially skewing the results and interpretations.

Despite the utility of modeling the relationship between connectomes, there is strong methodological disagreement in the literature as to how to extract the connectomes themselves from brain images^35–37^. Studies have shown that biases inherent in different processing choices, referred to here as “flavors”, can lead to different or even contradictory conclusions^38, 39^. Each connectome flavor may differ in how the brain is parcellated into regions, how potential noise, artifacts, or confounds are removed, and how the connectivity between regions is quantified, among other methodological choices. Furthermore, connectome data from different studies are often shared only in certain flavors, making comparisons between datasets challenging.

In this work, our fundamental assumption is that each set of choices in the imaging and processing pipelines provides a different view of the same underlying system. Drawing inspiration from recent advances in multi-view learning^40^, we develop and share a tool called “Krakencoder” that provides a way to combine these choices (i.e. connectome fusion), and thus create a more comprehensive representation of the brain’s connectivity. Deriving a unified latent representation from within-modality and/or across modality connectome estimates may allow reconciliation of differing views of the same underlying network in order to overcome limitations and combine benefits from various processing choices. Krakencoder is a novel “universal” brain connectivity encoding architecture and is capable of transforming connectivity estimates between parcellations, estimation techniques, and modalities with unprecedented precision. Importantly, the Krakencoder is able to preserve inter-individual differences in a way that is behaviorally and demographically relevant. This element is crucial if we are to use the Krakencoder to improve our understanding of how brain structure and function and their interplay maps to behavior, impairment, or recovery after disease or injury.

## 2 Results

### 2.1 Krakencoder model architecture and training

The Krakencoder architecture comprises a set of encoders and decoders that allow transformation of each connectivity type into every other connectivity type via a common low-dimensional latent representation, see **Fig. 1**. The Krakencoder was trained to optimize for both reconstruction accuracy and preservation of individual differences. The latter was achieved by enforcing latent-space representations of different connectivity types from the same subject to be close to one another and far from other subjects’ latent space connectome representations. The model was trained on resting-state FC and white-matter SC data from 683 healthy young adults (366 female, aged 28.6 ± 3.8) from the Human Connectome Project, validated during training on 79 held-out subjects (43 female, aged 29.1±3.8), and finally tested on 196 subjects (107 female, aged 29.1±3.6). Care was taken to ensure that no siblings spanned the divisions, as this data leakage can lead to over-optimistic model performance. Connectivity flavors include 3 parcellations (with 86, 268, and 439 regions), 3 common FC estimates (Pearson correlation, Pearson correlation following global signal regression, and regularized partial correlation), and 2 SC estimates (volume-normalized streamline counts from deterministic and probabilistic tractography), for a total of 15.

**Figure 1.**
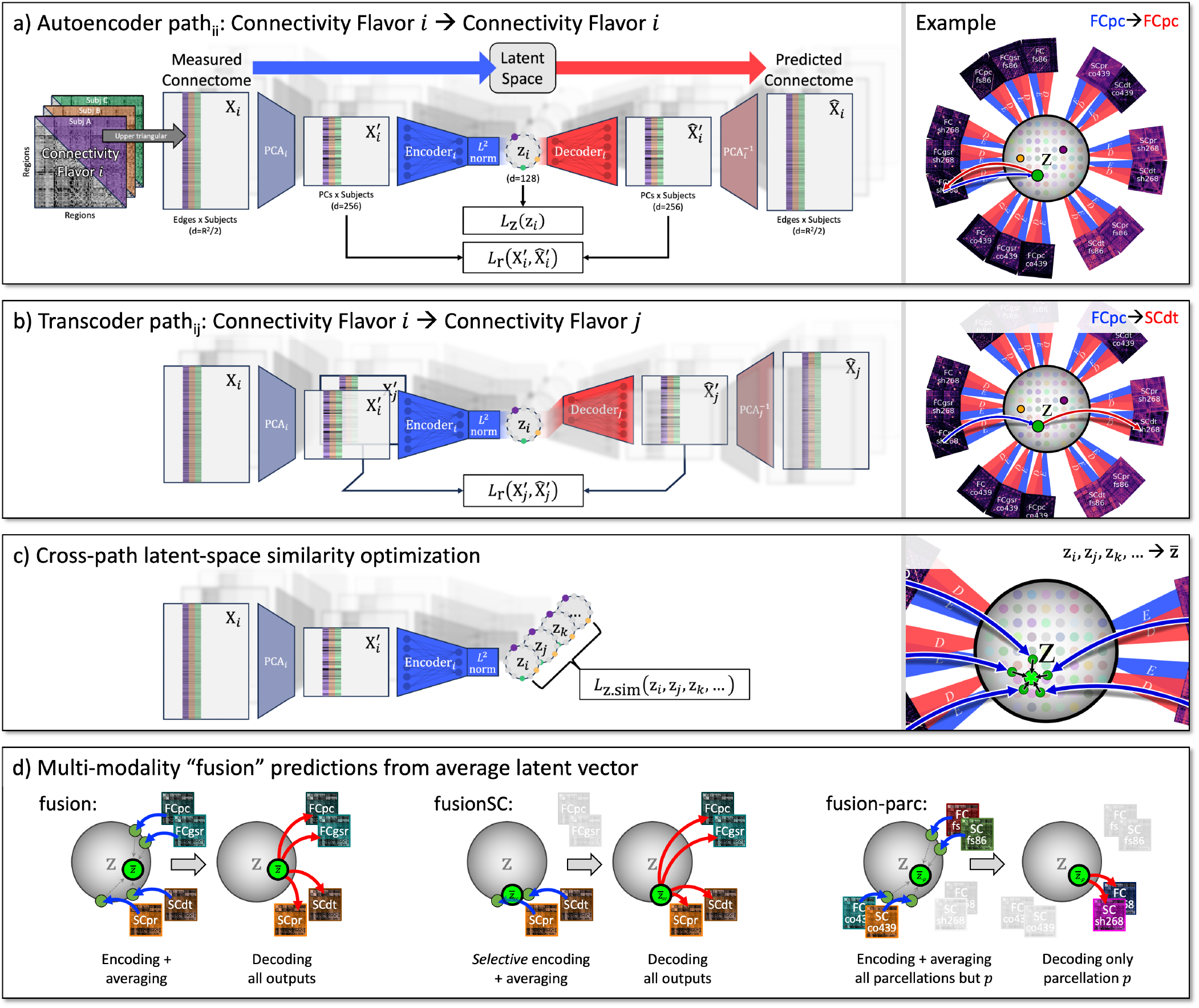
Key features of the Krakencoder architecture. **a**. Autoencoder for connectivity flavor *i* (i.e., *path*_*ii*_) begins by stacking the upper triangular portion of each subject’s connectivity matrix into input 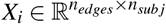. A precomputed, fixed PCA transformation normalizes the data and reduces dimensionality to 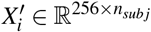, equalizing the size of disparate input flavors. A single fully-connected layer *Encoder*_*i*_ followed by *L*^2^ normalization transforms 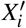 into a latent hypersphere surface 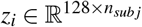. Batch-wise encoding loss *L*_*z*_(*z*_*i*_) controls inter-subject separation in latent space. A single fully-connected layer *Decoder*_*i*_ transforms *z*_*i*_ to 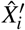, and batch-wise reconstruction loss 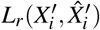 and *L*_*z*_(*z*_*i*_) are backpropagated to optimize *Encoder*_*i*_ and *Decoder*_*i*_. **b**. Transcoder *path*_*i j*_ converting input flavor *i* to output flavor *j* begins the same as *path*_*ii*_, transforming 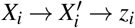, then *Decoder*_*j*_ transforms 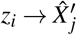, and reconstruction loss 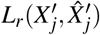 is backpropagated to optimize *Encoder*_*i*_ and *Decoder*_*j*_. **c**. Latent similarity loss *L*_*z*.*sim*_(*z*_*i*_, *z* _*j*_, *z*_*k*_, …) provides explicit control to ensure the latent representations of each subject are consistent across connectivity flavors. See **Table 1** for details about loss terms. **d**. Illustration of “fusion” predictions from averaged latent space vectors. For these predictions, we average the encoded latent vector from all input flavors (as in the “fusion” model), or a subset of input flavors (as in the “fusionSC” model which only averages latent vectors from SC inputs), and then decodes this average vector to all output flavors. The “fusion-parc” model demonstrates cross-parcellation prediction, by averaging only inputs from the *other* parcellations.

We create a separate encoder and decoder for each of the 15 connectivity flavors and then train the model to predict each flavor from each other flavor. There are 225 total training paths, including 15 autoencoders that map between the same flavor (See **Fig. 1a**) and 210 transcoders that predict one flavor from another (See **Fig. 1b**). Each encoder is a single fully-connected linear layer, followed by *L*^2^ normalization that constrains the latent representation to the surface of a 128-dimensional hypersphere. Each decoder is a single fully-connected linear layer. Dimensionality of the connectivity flavors varies between parcellations, which could lead to an imbalance in the training to favor higher dimension connectivity flavors. We balanced the contributions of each flavor by reducing each connectivity input to 256 dimensions using a precomputed PCA, derived using only training data to avoid leakage between train and test sets. Reconstruction loss for each training path includes both Pearson correlation and Euclidean distance. We include a contrastive component to the correlation and Euclidean loss functions, to explicitly preserve inter-subject variability of predicted connectomes. A contrastive loss term is applied to the latent representation as well to further promote inter-subject variability of the representations. The term “contrastive” here refers to a penalty that attempts to force the same subject’s reconstructed and latent space connectomes close to each other and far apart from other individuals’ representations. Finally, we include an intra-subject latent space consistency loss to maximize the similarity of the latent representations from each connectome flavor for a given subject (See **Fig. 1c**). This combination of loss functions allowed us to fine-tune the tradeoff between reconstruction accuracy and inter-subject variability (See **Fig. S2**), and to condition the latent space representation. See Methods for details on training procedure and loss functions.

**Table 1.** Loss function summary.

| Loss category |  | Equation | Description |
| --- | --- | --- | --- |
| Path loss | Reconstruction accuracy and inter-subject dispersion | $L_{r.corrI}(X'_i, \hat{X}'_i) = \ \text{corr}(X'_i, \hat{X}'_i) - I_{n_{subj} \times n_{subj}}\ _2$ | $[n_{subj} \times n_{subj}]$ correlation matrix of measured→predicted should be high along the diagonal and low off-diagonal |
| | | $L_{r.mse}(X'_i, \hat{X}'_i) = \ X'_i - \hat{X}'_i\ _2^2$ | MSE between measured→predicted should be low |
| | | $L_{r.marg}(X'_i, \hat{X}'_i) = -\min_{a,b \neq a} (\ X'_i(s_a) - \hat{X}'_i(s_b)\ _2)$ | Predicted $\hat{X}'$ for subject $s_a$ should be far from nearest measured $X'$ for $s_b \neq a$ |
| | Latent space inter-subject dispersion | $L_{z.corrI}(z_i) = \ \text{corr}(z_i, z_i) - I_{n_{subj} \times n_{subj}}\ _2$ | $[n_{subj} \times n_{subj}]$ correlation matrix of latent space vectors (for the same path) should be low off-diagonal |
| | | $L_{z.dist}(z_i) = -\sum_{a,b \neq a}^{n_{subj}} \ z_i(s_a) - z_i(s_b)\ _2$ | Mean inter-subject latent space distances should be high |
| | Combined path loss | $L_r(path_{ij}) = L_{r.corrI}(X'_j, \hat{X}'_j) + L_{r.marg}(X'_j, \hat{X}'_j) + w_{rm} * L_{r.mse}(X'_j, \hat{X}'_j)$ $(w_{rm} = 1000)$<br>$L_z(path_{ij}) = L_{z.corrI}(z_i) + L_{z.dist}(z_i)$<br>$L_{path} = L_r(path_{ij}) + w_z * L_z(path_{ij})$ $(w_z = 10)$ | |
| Latent space intra-subject consistency | | $L_{z.sim}(z) = \sum_{a=1}^{n_{subj}} (\sum_{i,j \neq i}^{n_{flav}} \ z_i(s_a) - z_j(s_a)\ _2^2)$ | Total MSE between all latent vectors from all flavors for each subject should be low |

During model inference, the Krakencoder predicts each of the 15 connectivity flavors from any other, transforming predictions that exist in the reduced 256-dimensional space to their higher native dimensions using the inverse of the precomputed PCA transforms. The model can also combine information from multiple connectomes by averaging their latent space representations to obtain a “fusion” representation; this fusion representation can then be decoded to predict all 15 connectome flavors. The encoding and averaging step in the fusion process can include all available connectome flavors, or a subset of flavors to, for instance, predict FC flavors from the fusion latent representation of all SC flavors or vice versa (“fusionSC” or “fusionFC”, **Fig. 1d**), or to predict one parcellation using inputs from the averaged latent representations of all other parcellations (“fusion-parc”, **Fig. 1d**). The latter fusion version is particularly informative if one is using the Krakencoder to translate connectome data to a desired atlas on which it had not been computed.

### 2.2 Measuring connectome prediction accuracy

A key consideration when assessing the performance of a connectome prediction model is the high degree of similarity of measured connectomes with the population mean. The Pearson correlation between connectomes of unrelated individuals can exceed 0.9, depending on the flavor, and a model that simply predicts the population mean for every subject can appear to have a very high prediction accuracy if it uses Pearson correlation as its metric (See **Fig. 2d**, “pop. mean” in bottom subpanel). One simple solution to this issue is to subtract the population mean calculated from training subjects before correlating measured and predicted connectomes; here we refer to this metric as avgcorr_demean_. However, this metric still cannot distinguish whether predictions faithfully capture individual variation, particularly when there is clustering within the population (see **Fig. S1b**). Our goal here is to create individualized connectome predictions to understand the sources and consequences of across-subject variability. Thus, we also consider the identifiability of our predictions: whether our measured connectome for a subject is closer to the predicted connectome for that subject than the predicted connectome of any other subject. The “top-1 accuracy” is a winner-take-all measure that reflects the fraction of subjects for which this is true. For a more continuous measure of identifiability, more suitable to assessing imperfect predictions, we use an “average rank percentile” measure (avgrank), which is the fractional rank of the similarity (Pearson correlation) of a subject’s measured and predicted connectomes compared to all other predicted connectomes, averaged across subjects (**Eq. 4**). Taken together, prediction accuracy and identifiability, as measured by avgcorr_demean_ and avgrank, convey a more complete picture of how well a model explains individuals’ connectomes.

**Figure 2.**
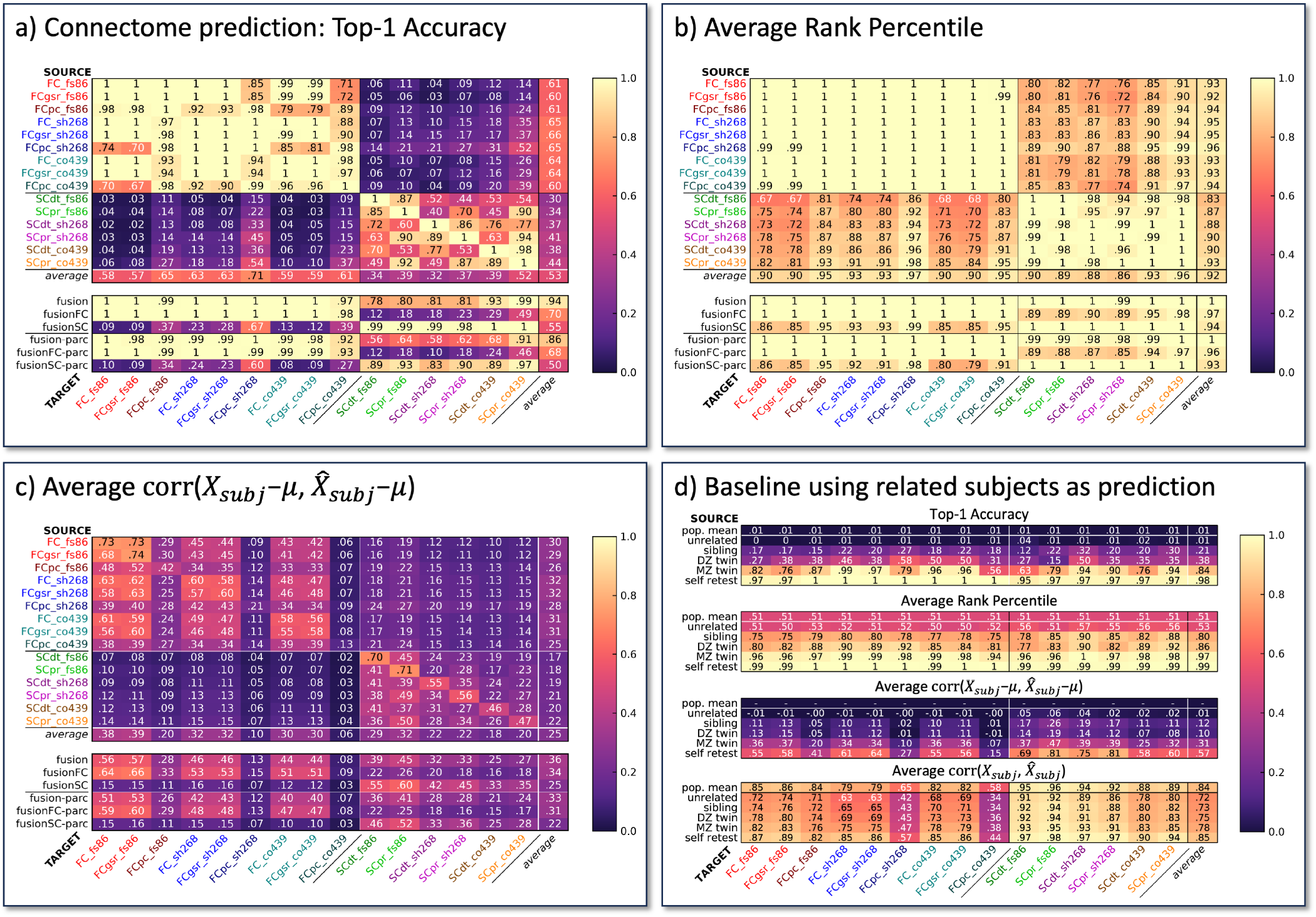
Connectome prediction performance for all input and output flavors on 196 held-out HCP-YA subjects. Heatmaps in **a-c** show connectome prediction performance from each *source* flavor (row) to each *target* flavor (column). Flavors include 3 parcellations (fs86, sh268, co439), 3 FC estimates (FC=Pearson, FCgsr=global signal regressed Pearson, and FCpc=regularized partial correlation), and 2 SC estimates (SCdt=deterministic tractography, SCpr=probabilistic tractography). The “*average*” row is the mean of performance metrics for all sources in the column above. The “fusion” row shows predictions from the average latent vector from all 15 connectome flavors (See **Fig. 1d**). “fusionFC” and “fusionFC” are predicted from the average of 9 FC or 6 SC latent vectors, respectively. “fusion-parc” predicts each parcellation from the average of all parcellations *except* itself. **a**. Top-1 accuracy measures the fraction of test subjects whose predicted connectome is closer to their own measured connectome than any other subject’s measured connectome. Chance level is 1/196=0.005. **b**. Average rank percentile measures the fraction of *other* subjects’ measured connectomes that are further from the target subject’s predicted connectome than its own measured connectome. This rank percentile is then averaged across subjects. Chance level is 0.5. **c**. Average correlation of each subject’s measured and predicted connectomes, after subtracting the population average (avgcorrdemean). **d**. For each subject *s*_*a*_, we select an age- and sex-matched individual *s*_*b*_ of a given relatedness (unrelated, sibling, DZ/MZ twin, or self retest) and consider the measured connectome from *s*_*b*_ as the prediction for *s*_*a*_. For each relatedness level, we compare the set of pseudo-predictions to measured connectomes using each performance metric, as though they were predictions from our model. Thus, we can observe the reliability and utility of each flavor and performance metric, and provide multiple baselines for evaluating model performance. We also show performance using the population average for each flavor (“pop. mean”) as pseudo-prediction, which for the avgcorr metric (bottom) exceeds test-retest, but for all other metrics is zero or chance.

In addition to these measures of prediction accuracy for the connectivity matrix values, we compute several common graph theoretical metrics to assess how faithfully the predicted connectomes capture the network properties of the measured connectomes. We use the Brain Connectivity Toolbox^41^ to compute average node strength, average betweenness centrality, characteristic path length, and modularity (Louvain community detection) for each measured and predicted connectome, and compute the Spearman rank correlation between the predicted and measured values across all test subjects for each metric.

### 2.3 Connectome prediction performance

The accuracy of the Krakencoder’s connectome predictions was assessed on 196 held-out test subjects via identifiability (top-1 accuracy and avgrank) and (de-meaned) correlation (avgcorr_demean_). The heatmaps in **Fig. 2a-c** illustrate the various metrics for every pair of the 15 connectome flavors, as well as “fusion” predictions decoded from the averaged latent vectors from groups of encoded connectome flavor inputs (See **Fig. 1d**). The “*average*” column and row show the average performance metric for each source and target flavor, respectively.

Top-1 identifiability (**Fig. 2a**) measures the fraction of subjects whose predicted connectome is more similar (higher Pearson correlation) to their measured connectome than any other subject’s measured connectome (random chance 1*/n*_*sub j*_ = 0.005). Average rank percentile (**Fig. 2b**) is the fractional rank of the similarity (Pearson correlation) of a subject’s predicted and measured connectomes compared to all other measured connectomes, averaged across subjects (random chance 0.5). The avgcorr_demean_ metric (**Fig. 2c**) is the average Pearson correlation between a subject’s predicted and measured connectomes, after subtracting the population average of the measured connectome flavor from the predicted and measures connectomes. The average rank percentile for FCs predicted from FCs of different parcellations or connectivity estimations (FC→FC) shows near-perfect identifiability (**Fig. 2b**, upper section, upper left block mean=1.0). Similarly, SC→ SC predictions have near-perfect identifiability (upper section, lower right block mean=0.99). Inter-modality predictions performed worse than within-modality, but still far exceeding chance, with the overall mean of SC→ FC (lower left block) and FC→SC (upper right block) average rank identifiability of 0.82 and 0.85, respectively. In general, we find better avgrank prediction identifiability coincides with connectome sparsity, with higher dimensional parcellations (Coco439) and more sparse FC estimates (FC_pcorr_) being more easily identifiable. The avgcorr_demean_ metric (**Fig. 2c**) shows an average of 0.38 for FC→FC, and an average of 0.35 for SC→SC. Inter-modality predictions are again more difficult, with average FC→SC=0.16, and SC→FC=0.09. Lower dimensional parcellations generally had higher avgcorr_demean_. Higher dimensional atlases are likely better at capturing inter-individual variability in the functional boundaries within an atlas and thus have higher identifiability, while lower dimensional atlases likely benefit from higher SNR within the measured regional SC and FC and thus have smaller reconstruction error when mapping between flavors.

Predictions of regularized partial correlation FC (FC_pcorr_) had the highest avgrank, but the lowest avgcorr_demean_ among FC flavors, which reflects the noise inherent in this estimate, even with regularization, but also an increased sparsity that drives identifiability. SC predicted from FC_pcorr_ yielded consistently better performance than other FC inputs, likely due to the increased sparsity and removal of non-specific correlations. The best single FC input for predicting all SC flavors was the Shen268 FC_pcorr_, which predicts Coco439 SC_pr_ with an avgrank of 0.99 and avgcorr_demean_ of 0.19. The best single SC input for predicting all FC flavors was the Coco439 SC_pr_.

The bottom subpanel of **Fig. 2a-c** shows connectome predictions using the “fusion” latent space average from multiple input flavors. The “fusion-parc” predictions exclude inputs of the same parcellation from the averaged latent vectors, and their relatively high values demonstrate the level of shared information across flavors within the Krakencoder’s unified latent space.

Finally, we show that the Krakencoder’s fusion prediction also preserves the network properties of the measured connectomes. **Fig. S6** shows the Spearman correlation between the measured connectomes’ network properties (mean node strength, mean betweenness centrality, characteristic path length and modularity) and the Krakencoder’s predicted connectomes’ network properties, across 196 held-out individuals in the test set. The three fusion representations are used as input and each of the 15 connectome flavors as output; correlations between measured network metrics from the various familial relatedness groups are shown as comparison. The Krakencoder’s predicted connectomes generally preserve the network properties of the measured connectomes as well as or better than the correlation between network properties extracted from identical twins (MZ) or two scans of the same individual (self-retest).

### 2.4 Connectome similarity of related individuals

The “fusionSC-parc” and “fusionFC-parc” rows in **Fig. 2a-c** represent the accuracy of the Krakencoder when predicting an SC or FC connectome flavor from all other SC or FC flavors (excluding those from the predicted parcellation). If we compare this directly to **Fig. 2d**, we can see how the Krakencoder’s predicted connectome accuracy compares to several baseline estimates: measured connectomes from age- and sex-matched individuals who are unrelated, family members (non-twin siblings, dizygotic [DZ] and monozygotic [MZ] twins), retest sessions from the same individual (self retest), or the population mean connectome. For instance, the “self retest” row in the **Fig. 2d** heatmaps can be considered a measurement reliability ceiling, while the “unrelated” row shows the performance that can be explained by age- and sex-matching alone. Within a modality, top-1 accuracies (ranging from 0.93-1 for “fusionFC-parc”→FC and 0.83-0.97 for “fusionSC-parc”→SC) and average rank percentiles (all equal to 1 for the “fusionFC/SC-parc”) in **Fig. 2a** and **Fig. 2b** are similar to the self retest identifiability (when connectomes derived from imaging the same person twice are used to perform the identification) and better than the identifiability of individuals who are genetically identical (MZ twins). Similarly, the diagonal entries in **Fig. 2c**, representing the autoencoder avgcorr_demean_ for each flavor, range from 0.13-0.74, which is comparable to the “self retest” noise ceiling (0.15-0.81), and are larger than the “MZ twin” row (0.07-0.47, a 40-110% increase). This indicates that the autoencoders with a 128-dimensional bottleneck preserve more inter-subject connectome variability than can be explained by identical genetics, and about the same amount of variability as expected when measuring the connectome from the same individual twice. The “fusionSC” and “fusionFC” rows in **Fig. 2c** represent the avgcorr_demean_ for the unified SC (or FC) → FC (or SC) connectome predictions, with values ranging from 0.03 to 0.16 for SC→FC and 0.16-0.26 depending on the flavor. These exceed the similarity observed in the “sibling” or “DZ twin” row from the avgcorr_demean_ subpanel in **Fig. 2d**, indicating that predicting FC from our unified SC representation preserves more intersubject FC variability than can be explained by age, sex, and 50% genetic similarity.

### 2.5 Spatial variation in edge prediction accuracy

The primary performance metrics assess global connectome similarity. We further assessed the spatial variability of prediction accuracy to determine whether certain brain regions are more accurately predicted than others. For fusion-parc, fusionFC-parc, and fusionSC-parc (fusion predictions made without the parcellation being predicted), the prediction accuracy for each edge is the Pearson correlation between the predicted and measured values for the 196 held-out test subjects. To simplify visualization and enable comparison across varied size parcellations, each region is assigned to one of 7 cortical resting state networks^42^ or subcortical and cerebellar networks. The edge-wise accuracies (region×region) are then grouped and averaged into a 9× 9 network representation for each connectome flavor. In the top block of **Fig. S3a**, the network-averaged edge prediction accuracy is presented for each of the fusion types as input (rows) and each connectivity flavor as output (columns). The bottom block shows the baseline network-averaged edge similarity comparing measured connectomes of unrelated age- and sex-matched subjects and related individuals. The figure demonstrates that the Krakencoder’s prediction accuracy is relatively uniform throughout the cortex and subcortex. Networks with lower Krakencoder predictability (e.g., the limbic network), also show low self-retest measurement reliability (bottom row). In **Fig. S3b**, each network × network heatmap in **Fig. S3a** is correlated with the corresponding self-retest measurement reliability (bottom row), to quantify the consistency of the spatial pattern of edge-wise accuracies with the reliability of the edge-wise measurements. We see that the spatial pattern of the edge-wise accuracy of the Krakencoder’s fused intra- and inter-modality predictions is as similar to the spatial pattern of the edge-wise self-retest measurement reliability as the MZ twins’ spatial patterns of edge-wise reliability. **Fig. S4** summarizes the edge-wise prediction accuracy at a region level for each connectivity flavor in its own parcellation, showing a similar agreement between prediction and measurement reliability.

We observe that higher level association cortex (e.g., default mode) tends to have higher SC→FC prediction accuracy than sensory-motor cortex (e.g., visual). In **Fig. S5a**, we further quantify this relationship by correlating regional prediction accuracy with an estimate of the cortical hierarchy gradient derived from functional connectivity^43^, and find a significant positive correlation (Spearman *r*=0.249, *p*_*spin*_=0.019, 10000 rotations^44^). Thus, our SC→FC predictions explain more of the population variability in higher level cortex than in lower. Conversely, if we correlate corresponding rows in the square measured and predicted connectome matrices, we find that predicted connections to high-level cortex are less coupled to their measured rows than low-level cortex (Spearman *r*=-0.191, *p*_*spin*_=0.116; see **Fig. S5b**). This latter relationship is consistent with previous work on structure-function coupling^16, 19^, but primarily reflects the relationship between population averaged SC and FC rather than predicting inter-individual variability.

### 2.6 Demographic and behavioral prediction from the Krakencoder’s latent space representations

We next tested if the Krakencoder latent space preserves inter-subject variation related to family structure as well as demographic and behavioral features. **Fig. 3a** shows the inter-subject similarity of latent space representations for pairs of age- and sex-matched unrelated individuals, non-twin siblings, DZ and MZ twins, and self-retest, along with inter-subject similarity of observed connectomes. By comparing ROC separability of each pair of violin plots, we see that separation is greater for the Krakencoder latent space than for the observed connectomes, with statistical significance for unrelated vs sibling, DZ vs MZ twins, and MZ vs unrelated (*p*_*perm*_ *<* .05, 10000 permutations, FDR-corrected).

**Figure 3.**
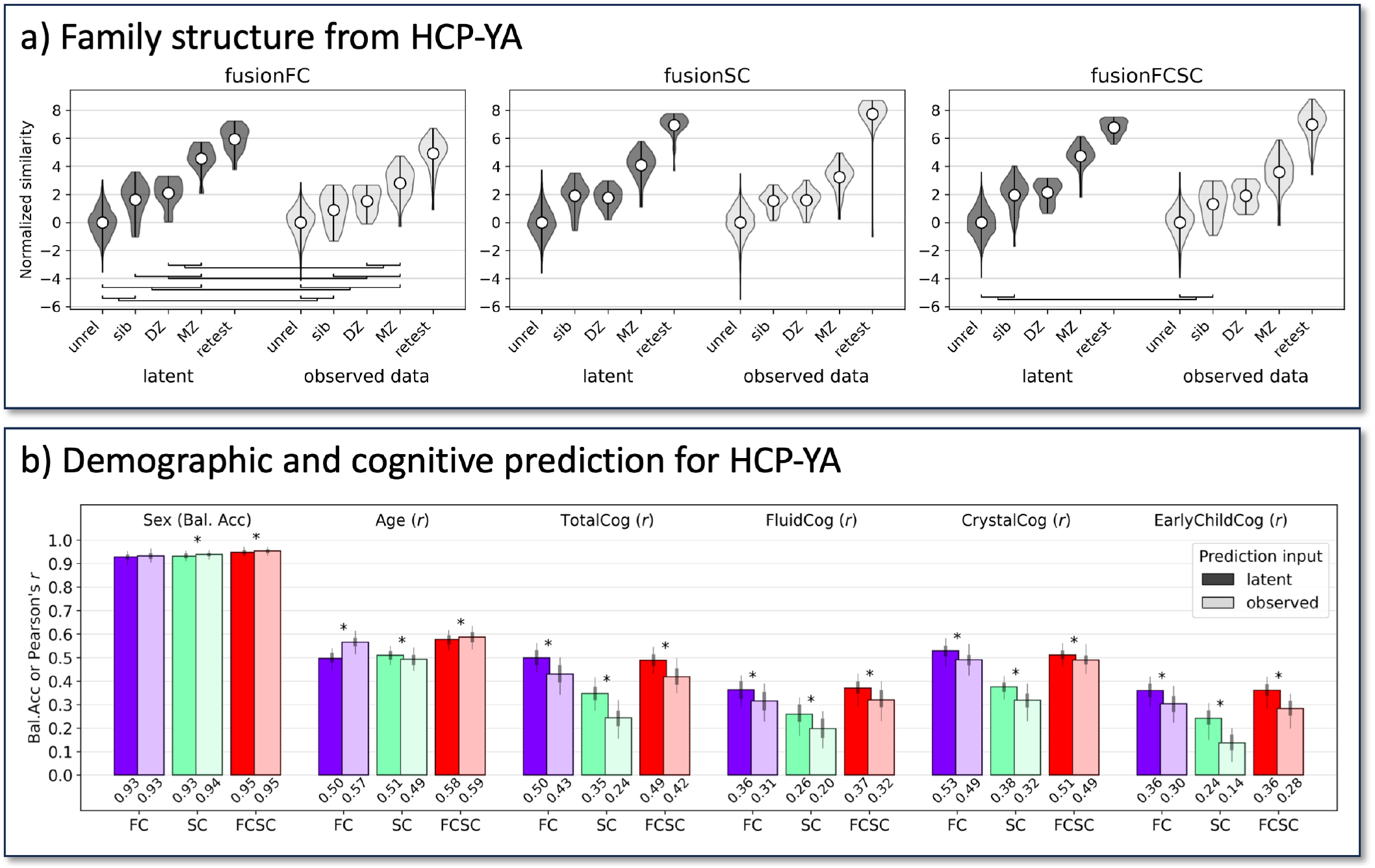
Familial connectome similarity and demographic prediction for held-out HCP-YA subjects. **a**. Inter-subject similarity of latent space representations and observed SC/FC data, stratified by family relationship. Each violin plot shows the distribution of inter-subject similarity for age- and sex-matched pairs for each familial relationship, as well as unrelated pairs and self-retest. Due to the high baseline similarity in observed data, latent and observed data similarity measures were independently z-scored by the mean and standard deviation of the similarity of unrelated subjects for visual comparison. Separability of each pair of violins within a modality was computed as the area under the receiver operating characteristic (ROC) curve. Separability estimates for a given relationship pair from the latent space and observed data were statistically compared using a permutation test, where bars show family-level pairs where latent space separability exceeds observed data separability (*p*_*perm*_ *<* .05, 10000 permutations, FDR-corrected). **b**. Prediction accuracy of demographic and cognitive metrics from different subsets of the Krakencoder’s latent space and observed SC and FC data. Variability of prediction accuracy was assessed through bootstrap resampling (N=100). *= significant difference between prediction accuracy from latent space and observed data. (*p*_*perm*_ *<* 10^−3^, 10000 permutations, FDR-corrected).

To demonstrate the preservation of demographically- and behaviorally-relevant features in the Krakencoder’s latent space, we trained linear models to predict individuals’ age, sex, and cognitive scores (“total”, “fluid”, “crystallized”, and “early childhood” composite scores from the NIH toolbox) from the Krakencoder latent space, and compared them to the same predictions based on observed connectome data. Linear kernel ridge regression models for each variable were built from the mean latent vectors (fusionFC, fusionSC, fusionFCSC, each of which is *n*_*sub j*_ × 128) or from the average inter-subject cosine similarity of observed data (i.e. the average of 9 FC, 6 SC, or 15 SC+FC *n*_*sub j*_ × *n*_*sub j*_ matrices). **Fig. 3b** shows Krakencoder and observed connectome data predict sex with a similar balanced accuracy of ≥ 0.93. The Krakencoder’s latent space predicts age with Pearson *r* ≥ 0.5, (fusionFC *r*=0.50, MAE=2.35; fusionSC *r*=0.51, MAE=2.57; fusionFCSC *r*=0.58, MAE=2.33), similar to the predictions from observed data (observed FC *r*=0.57, MAE=2.31; observed SC *r*=0.49, MAE=2.58; observed FC+SC *r*=0.59, MAE=2.33). For the age predictions, fusionSC outperformed observed SC and observed FC outperformed fusionFC (*p*_*perm*_ *<* 10^−3^, FDR-corrected), but the observed data and fusion SCFC models performed similarly. Interestingly, the Krakencoder’s latent space predictions significantly outperformed the observed data model predictions for predicting all cognitive composite scores. See Methods for details on inter-subject similarity calculations and model fitting.

### 2.7 Sensitivity analyses

Sensitivity analyses were performed to determine which brain areas’ connections played the largest role in i) the accuracy of the Krakencoder’s connectome mapper, and ii) the accuracy of the models predicting age, sex, or total cognition from the Krakencoder’s latent space. These analyses demonstrate how the Krakencoder may be used to understand the biological underpinnings of, for example, SC to FC relationships or what brain networks may play the largest role in predicting cognition. For simplicity, regions were grouped into 7 cortical resting state networks^42^ or subcortical and cerebellar networks. To perform the sensitivity analyses, every connectome edge was set to the population mean of the training set, except for the columns/rows corresponding to the regions within the network of interest, which were left untouched, see **Fig. 4a**. The fusion latent space representations (fusion, fusionSC, and fusionFC) were re-calculated for each individual and used to predict i) each connectome flavor and ii) age, sex and total cognition using re-fitted KRR models.

**Figure 4.**
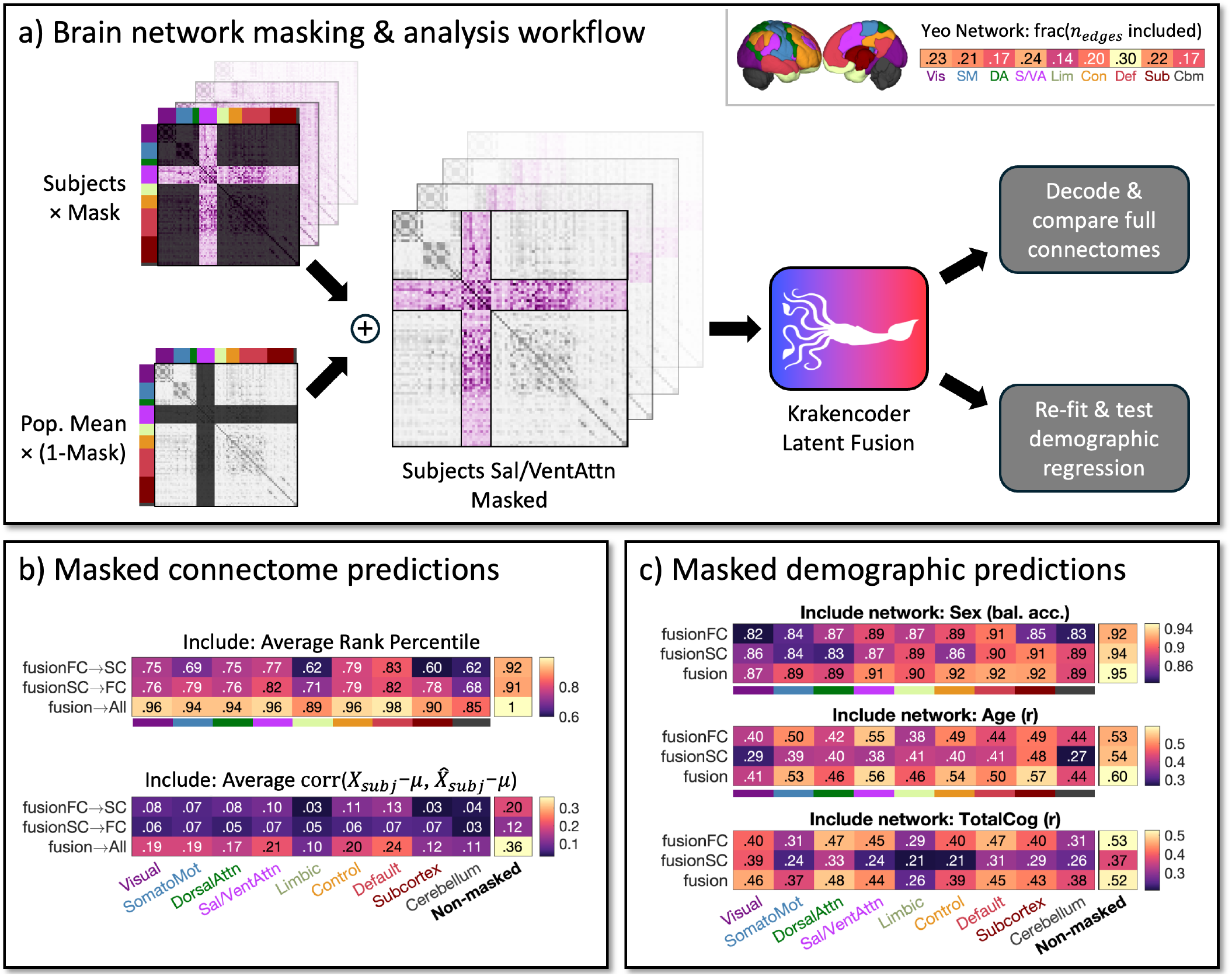
Spatial input sensitivity analysis of Krakencoder predictions. **a**. We estimate the sensitivity of the Krakencoder’s prediction accuracy to regions within a given network by replacing all connectome edge values not in that network with the population mean and feeding these masked connectomes into the connectome and demographic predictions. The accuracy after masking reveals the amount of information that particular network’s regional connections have in the mapping and demographic/behavioral predictions. **b**. Connectome prediction identifiability (avgrank, top) and reconstruction accuracy (avgcorr_demean_, bottom) using only the connections to/from regions within each brain network shows how much information the model is utilizing from each network when predicting whole-brain connectomes. The “non-masked” column on the right shows performance using original connectomes without masking. **c**. Similarly, predicting subject sex (top), age (middle), and total cognition (bottom) from masked input data shows the relative amount of information the connections to/from regions in each network contain about those demographic and cognitive features.

**Fig. 4b** shows the mean avgrank and avgcorr_demean_ when only individual-level values of those networks’ con-nections’ were retained. The accuracy metrics for the fully intact, non-masked connectomes are shown on the right; values that are lower indicate that network’s connections do not contribute as much to the accuracy of the Krakencoder’s mapping. The top row shows the mean avgrank over the SC flavors predicted from fusionFC, the second row shows the mean avgrank over the FC flavors predicted from fusionSC and the third row shows the mean avgrank across all flavors of predicted SC and FC. We see that default mode network connections appear important for mapping accuracy, particularly for the fusion →all and fusionFC→SC. Connections to/from regions in the somatomotor network and subcortex seem to be more informative for fusionSC→FC than for fusionFC → SC.

In **Fig. 4c**, we see the largest retention in accuracy for predicting sex when functional and structural connections to/from regions in the default mode network and structural connections to/from the subcortex are retained. Age was best predicted using functional connections to/from the salience/ventral attention network and structural connections to/from the subcortex. Total cognition was best predicted by functional connections to/from regions in default mode and dorsal attention networks, and by structural connections to/from regions in the visual network.

### 2.8 Krakencoder performance on out-of-sample, out-of-distribution data

To assess generalizability of the Krakencoder to new datasets, we applied our frozen, pretrained Krakencoder model to data from the multi-site HCP Lifespan study^45–47^, including both the Development cohort (HCP-D) and the Aging cohort (HCP-A). These two studies differ from the original HCP young adult study in both subject demographics (ages 8-21 for HCP-D and 36-100+ for HCP-A vs 22-37 for HCP-YA) and acquisition parameters, with voxel sizes and scan durations more similar to those used in typical neuroimaging studies (see Methods for more details). To account for domain shift, we linearly mapped the population mean of each connectome flavor to the population mean of the HCP-YA training subjects before applying the HCP-YA derived PCA dimensionality reduction and Krakencoder latent space projection. Despite the demographic and acquisition differences, the Krakencoder still predicts connectome flavors with high accuracy and precision. **Fig. 5a-d** shows avgrank and avgcorr_demean_ for the HCP-D (age 8-21) and HCP-A (age 36-100+) cohorts. Within-modality connectome prediction identifiability remained high for both HCP-D and HCP-A (avgrank > 0.96). Inter-modal prediction identifiability was reduced 6-12% compared to held-out HCP-YA subjects, though still well above chance (avgrank > 0.75). Intra-modal avgcorr_demean_ was reduced by 10-26%, while inter-modal avgcorr_demean_ was 30-50% lower than held-out HCP-YA. Thus, despite significant differences in age and acquisition reducing prediction accuracy, the Krakencoder model still performs well above chance and preserves inter-individual differences when mapping between connectome flavors.

**Figure 5.**
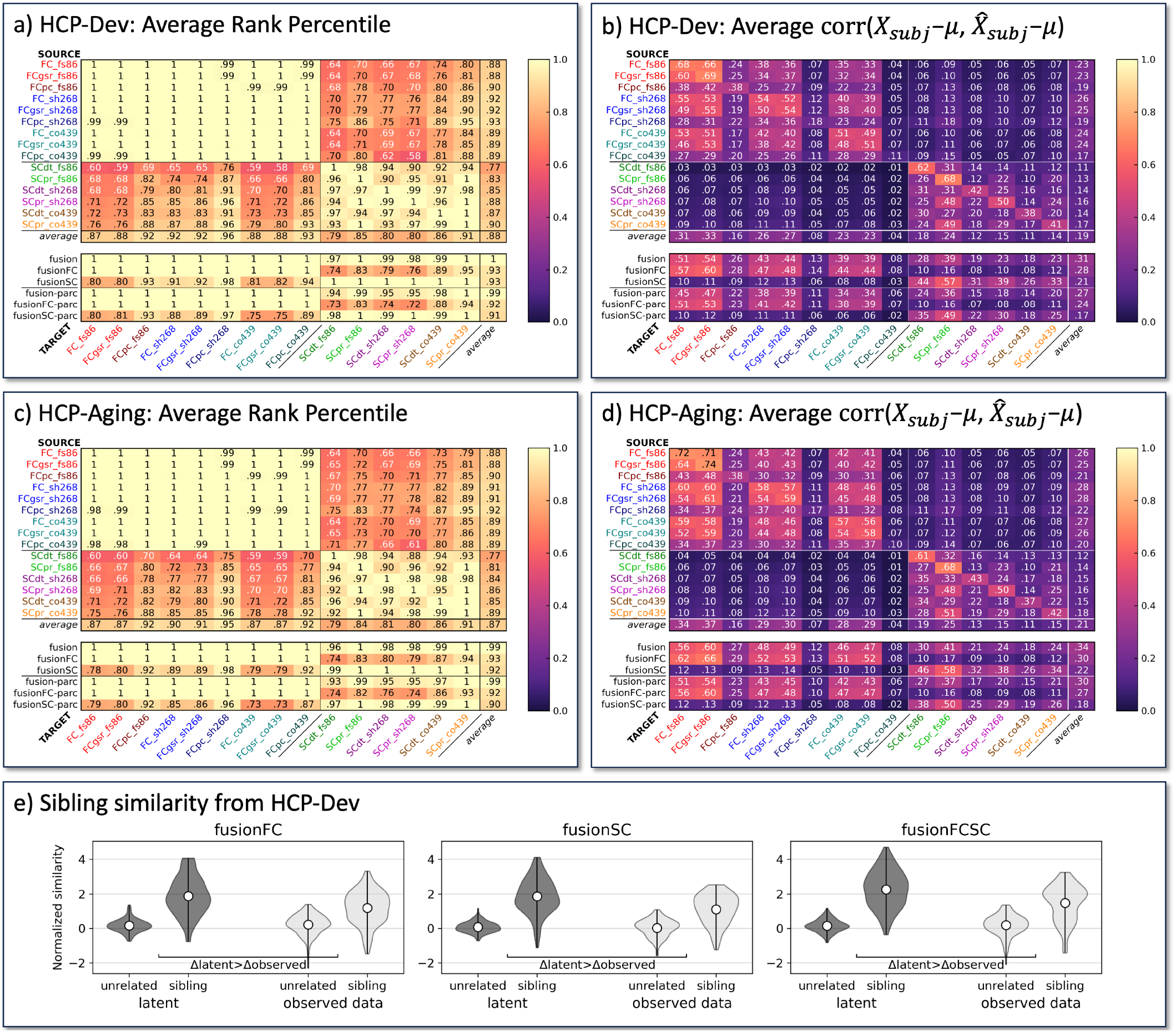
Out-of-sample performance of the Krakencoder on individuals from across the lifespan. Heatmaps in **a-b** show connectome prediction performance for subjects from the HCP-Development study (ages 8-21) and heatmaps in **c-d** show connectome prediction performance for subjects from the HCP-Aging study (ages 36-100). The Krakencoder model applied was trained on HCP-YA data (ages 22-37) and frozen; no re-training or fine-tuning was performed. **a**,**c**. Average rank percentile (**Fig. 2b**) is the fractional rank of the similarity (Pearson correlation) of a subject’s predicted and measured connectomes compared to all other measured connectomes, averaged across subjects (random chance 0.5). **b**,**d**. Average correlation of each subject’s measured and predicted connectomes, after subtracting the population average. **e**. Connectomes from sibling pairs in HCP-D are more similar than age- and sex-matched unrelated pairs, with latent space similarity significantly more separable than observed data. Each violin plot shows the distribution of inter-subject similarity for sibling pairs or pairs of unrelated individuals with matching age and sex disparity. As in **Fig. 3a**, latent and observed connectome similarity were independently normalized for visualization. Separability was measured by ROC analysis, and significance was assessed using permutation testing. For all three fusion types (fusionFC, fusionSC, fusionFCSC), siblings were more similar in the Krakencoder’s latent space than in the observed data (*p*_*perm*_ *<* 10^−3^).

The HCP-D dataset contains 76 sibling pairs (50 same-sex, 26 opposite sex, Δage 0-9yr), so we performed a familial similarity analysis similar to that for HCP-YA. In **Fig. 5e**, we show that siblings were more separable from age- and sex-matched individuals in latent space than in the observed data for all three fusion types (fusionFC, fusionSC, fusionFCSC) (*p*_*perm*_ *<* 10^−3^, FDR-corrected). Finally, we performed a regression analysis to predict age, sex, and cognition in the HCP-D and HCP-A datasets (See **Fig. S7**). New predictive models using the same approach as the HCP-YA predictions were fit and cross-validated for each of the two new datasets. As with HCP-YA, sex was predicted very accurately by both latent and observed data models (balanced held-out test accuracy = 0.86-0.95). Age was predicted with much higher accuracy than in HCP-YA (HCP-D *r*=0.84-0.95, MAE=1.45-1.72; HCP-A *r*=0.76-0.85, MAE=6.13-6.52 on held-out test data), likely due the much larger age range and larger age-related changes during development and aging. Cognitive score prediction performance for the HCP-A data was similar to or better than HCP-YA, perhaps due to larger age-related cognitive effects, but performed considerably worse for the HCP-D dataset. This decrease in predictive ability was especially pronounced when predicting cognition from SC, where accuracy (Pearson’s *r*) from latent and observed data models was not significantly greater than 0. For all other predictions, observed data models performed significantly better than those on the Krakencoder latent space (*p*_*perm*_ *<* 10^−3^, FDR-corrected), though the Krakencoder predictions remained well above chance.

We aim to further test Krakencoder’s biological plausibility by applying it to connectomic data from individuals with pathological changes in the brain’s structural and functional connections. We extracted SC and FC from 100 individuals with multiple sclerosis (age: 22-71, 45.5 ± 11.4 years; 66% female; disease duration: 12.97±8.07 years), an autoimmune disease that results in lesions primarily in the brain’s white matter^48^. This disease has also been associated with changes in SC and FC that are related to individuals’ disability levels^49, 50^. The fMRI and dMRI acquisitions for these patients had significantly lower spatial, temporal, and angular resolution than the HCP-YA or HCP-Lifespan data (see **Table S1**). We fed the observed, pathological SC and FC into the corresponding arms of the frozen Krakencoder to obtain a fusionSC and fusionFC, from which we predicted the various FC or SC flavors. We found that the average rank identifiability of the Krakencoder’s predicted FC (from the fusionSC) was up to 71%, which was higher than the identifiability of the observed SC to observed FC (calculated within the same atlas), which was at most 54%, see **Fig. S8**. Mapping the fusionFC, calculated from fMRI in the MS patients, to the various SC flavors using the Krakencoder had around the same identifiability (up to 63%) as the mapping between observed FC to SC. These results demonstrate that even though the Krakencoder was trained on young, healthy individuals, it can still represent with good fidelity the relationship between connectomes in individuals with pathology. Importantly, it may indeed better capture the mapping from SC to FC than the observed connectome data itself. This provides evidence of the biological plausibility of the Krakencoder’s architecture and hints at its possible clinical significance in capturing low-dimensional representations of disease pathology.

### 2.9 Comparison with existing techniques

While predicting FC from SC is not the only application of our model, this route of prediction is among the most commonly studied problems in brain connectivity mapping. To compare our current Krakencoder model with this previous literature, we gathered a few recent studies that used different approaches to predict FC from SC. The first is a deep neural network proposed by Sarwar et. al^24^ (“deepnet”) and the second is a graph neural network from Neudorf et. al^25^ (“graphnet”). The Krakencoder model incorporates 6 flavors of SC (2 per parcellation) and 9 flavors of FC (3 per parcellation). These two previous SC-FC prediction frameworks are designed to predict a single flavor of FC from SC. Thus, we trained 12 separate models for each of the two frameworks for all combinations of SC_dt_/SC_pr_ and FC/FC_pcorr_. FC_gsr_ was excluded due to computational constraints and relative redundancy with FC. By both avgrank and avgcorr_demean_ metrics, the predictions from our model outperform both “deepnet” and “graphnet” models across all connectome flavors and parcellations (See **Fig. S9**). Specifically, the Krakencoder avgrank was 17-86% higher (average 42%) than “deepnet” or “graphnet” models for the same single SC inputs, and 18-90% higher (average 54%) using the fusionSC input to predict each FC flavor. Krakencoder avgcorr_demean_ was 1-890% higher (average 335%) than either model using single SC inputs, and 25-980% higher (average 450%) using fusionSC. Similarly, the accuracy of network properties for FC predicted by Krakencoder exceeded that from either alternative SC to FC framework. These particular results highlight the special emphasis of the Krakencoder on preservation of individual differences when mapping from SC to FC.

A note on computational efficiency: training all 12 “deepnet” models required approximately 140 GPU hours on an Nvidia A100 (4×5 hours for FS86, 4×10 hours for Shen268, and 4×20 hours for Coco439), and all 12 “graphnet” models required 1200 GPU hours (4×50 hours for FS86, 4×100 hours for Shen268, 4×150 hours for Coco439). For comparison, the Krakencoder required only 30 GPU hours to train a model for all 15×15=225 prediction paths simultaneously. When applying the Krakencoder model to new connectome data, the computational requirements are minimal and can even be done using free cloud-based platforms like Google Colab.

### 2.10 Extending the Krakencoder to new connectivity flavors

To broaden the applicability of our model, we demonstrate a means to extend the pre-trained Krakencoder model beyond the 15 flavors shown here. As depicted in **Fig. S10a**, given a new connectivity flavor computed on the same training subjects as the pre-trained model, we train a new autoencoder with an additional loss term to force the new latent representation into alignment with the pre-computed fusion latent representation. To illustrate this capability, we extended our model to new data from a previously unseen parcellation with bandpass-filtered time series for FC (original flavors use a high-pass filter) and un-normalized streamline count for SC (original flavors use pairwise volume normalization). As seen in **Fig. S10b**, prediction identifiability and reconstruction accuracy are comparable to the original 15 flavors. Furthermore, unlike training the original 225 path model, which takes up to 30 hours on a GPU, adding each new flavor takes only 1-2 minutes without a GPU. In this example, we used the original 683 HCP-YA training subjects, but this model-extension technique can use any population for which have already computed the Krakencoder latent representation.

## 3 Discussion

Here, we present the Krakencoder, a joint connectome encoder/decoder tool, capable of simultaneously, bidirectionally translating between structural and functional connectivity, and connectomes from different atlases and processing choices. The Krakencoder’s predicted connectomes match measured connectomes with an unprecedented level of accuracy and individual-level identifiability, while also preserving their network properties. The Krakencoder’s across modality mapping (from SC to FC and vice versa), which is arguably the most commonly performed mapping, had identifiability that was 42-54% higher than existing deep learning models. In addition to translating between connectome flavors, the Krakencoder can fuse connectome representations to create a single low-dimensional representation of all measured connectome flavors, each of which provide a different view of the true underlying connectome. The latent space of the Krakencoder appears to i) better reflect familial relatedness (compared to observed connectome data), ii) preserve age- and sex-relevant information and iii) enhance cognition-relevant information (again, compared to the observed connectome data). Finally, we demonstrated that the Krakencoder can be applied without any retraining to completely new, out-of-age-distribution lifespan data and still preserve inter-individual differences in the connectome predictions and familial relatedness in the fused latent space representations. Overall, we believe the Krakencoder has several novel aspects and applications, and is a significant leap forward in quantifying the relationship between multi-modal brain connectomes and how these relationships map to behavior.

Previous work in mapping between connectomes, mostly predicting FC from SC, has largely focused on optimizing accuracy in the model predictions by maximizing the correlation between predicted and measured FC. However, we argue here that over-emphasis on this metric can result in a model that predicts FC that is close to the population mean and/or does not preserve individual differences. The “deepnet” model did incorporate into its loss function a term that aimed to reduce inter-subject similarity of predicted FC^24^, however it appears that this only pushed the individual FC predictions away from one another and not toward the particular individual’s measured connectomes as its identifiability was just above chance. The Krakencoder’s connectome predictions, on the other hand, achieve perfect average rank identifiability within the same connectome modality, and consistently high average rank identifiability predicting FC from SC. Other modeling work has revealed that regional SC-FC coupling follows a functional gradient wherein the coupling is highest in visual and somatomotor networks and lowest in higher-order association networks^16, 19^. The Krakencoder predictions of FC from fusionSC follow this trend where the regional, row-wise correlation of observed FC and the Krakencoder’s predicted FC from fusionSC appears to be linearly decreasing with increasing regional hierarchy (see **Fig. S5**). This metric of row-wise correlation is very different from the across-subject, network-level correlations we are illustrating in **Fig. S3** that appear to be inversely correlated with regional hierarchy. As the Krakencoder architecture predicts each edge from all other regions and edges, rather than a single row, the edge-wise accuracy appears to be more aligned with measurement reliability than regional coupling. (see **Fig. S3** and **Fig. S4**).

The Krakencoder has many novel aspects, including its ability to translate between connectome flavors without requiring raw data. This is useful when researchers only have access to a connectome from a particular pipeline or atlas and need it in another. One previously developed tool called CAROT does allows translation of a connectome from a different atlas into another without requiring the raw data^30^; however, this tool does not allow translating connectomes between different processing pipelines or mapping between SC and FC. There is not one FC or SC processing pipeline that has been agreed upon by the field; many researchers chose one pipeline and replicate on others. The ability of the Krakencoder to predict other flavors within the same modality suggests a common, latent connectomic backbone that gives rise to each FC or SC flavor when a given parcellation or connectome estimation strategy is employed. The Krakencoder’s fusion functionality removes the burden of having to choose one pipeline or replicate findings on several different pipelines and attempt to make general conclusions. More than that, we believe the fusion of various snapshots of the underlying connectome increases the SNR of the connectome representations and perhaps becomes more than the sum of its parts. This latter statement is supported by the finding that models predicting cognition from the fused latent space representations outperformed models based on the observed connectome data; age and sex predictions were similar between the models but these are generally easier to predict compared to cognitive scores. The Krakencoder’s SNR boosting effect will be especially useful when the tool is applied to connectomes extracted from lower-quality data, which is often the case in clinical populations. Here, using lower-quality MRI data from individuals with multiple sclerosis, we demonstrate that the fusionSC to FC mapping has better identifiability (and the fusionFC to SC mapping similar identifiability) than the observed connectome data. The Krakencoder is robust when applied to completely new, out-of-distribution data from individuals across the lifespan (8-100+) and from adults with pathological changes to their connectomes (individuals with multiple sclerosis, or MS). The frozen model (trained on young adult data) still retained a high level of identifiability and explained variance when mapping between connectomes from the HCP-D, HCP-A, and MS datasets. The Krakencoder appears somewhat impervious to the sweeping connectomic changes that occur throughout human development, aging, and, to a certain extent, disease processes^51, 52^. The drop in demographic predictions of the latent space for the HCP-D, HCP-A, and MS data may mean there may be a benefit of the Krakencoder from further training (perhaps fine tuning) using lifespan and /or disease data.

Future work will apply the Krakencoder to predict connectomes under other clinical conditions, and further explore the relationship between the common latent representation and factors of neuroscientific or clinical significance. We will also create a fine-tuning procedure to further increase the robustness of the Krakencoder when applied to novel datasets in order to promote its wider applicability, particularly in datasets that contain pathology like stroke and Alzheimer’s disease. The Krakencoder could also be adapted to have encoding and decoding arms for data from other modalities (MEG or EEG) or even to explicitly capture demographic/behavioral information so that the latent space may be used more effectively for brain-behavior mapping. While we purposefully chose atlases that were derived in varied ways, i.e. using anatomical vs functional information, there still may be disagreement in the labeling of regions across subjects. However, by translating between and fusing SC and FC derived from all atlases, the Krakencoder attempts to identify a latent organizational representation that underlies those observed in different overlapping parcellation schemes. In the context of SC-FC coupling, we do not think these atlas choices prevent its accurate evaluation within an individual, since we are using the same parcellation for both SC and FC. Finally, while this first proof-of-concept work focused on applying the Krakencoder to static FC, we know that brain activity is flexible and adapts to task demands, varied states, and pharmacological manipulations. Further work will investigate adapting the Krakencoder to better reflect dynamic changes that occur in brain networks.

## 4 Methods

### 4.1 Data description

The data for this study come from the original Human Connectome Project (HCP)^53^ and the HCP Lifespan studies^45–47^, which together includes high-resolution 3T MRI data for 2490 healthy participants aged 8-100+. Primary model training and testing was performed using data from 958 subjects from the original “young adult” HCP study (HCP-YA), aged 22-37, which includes monozygotic and dizygotic twin pairs as well as non-twin siblings. For benchmarking performance, we include a subset of 40 HCP-YA subjects who returned for a complete “retest” session 1-11 months after their initial session. The HCP-Development (HCP-D, 608 subjects, aged 8-21, including 76 pairs of non-twin siblings) and HCP-Aging (HCP-A, 716 subjects aged 36-100) cohorts from HCP Lifespan were used to evaluate our pre-trained model on out-of-sample, out-of-distribution data. HCP Lifespan data differs from HCP-YA in both age range and acquisition parameters, with Lifespan acquiring less than half as much fMRI and dMRI data as HCP-YA. HCP Lifespan studies were also collected across four different sites using standard MRI scanners while the HCP-YA data were all acquired on a custom MRI scanner at a single site. See **Table S1** for demographic and acquisition details. For all three datasets, we excluded subjects with incomplete resting-state fMRI or diffusion MRI, and subjects with known acquisition or QC issues.

### 4.2 Construction of Functional Connectomes

All resting-state fMRI timeseries were preprocessed by HCP using the Minimal Preprocessing Pipeline^54^, which includes motion correction, susceptibility distortion correction, coregistration to anatomy and nonlinear template space (FSL MNI152 template, 6th generation), as well as automated denoising using ICA-FIX^55^. We used a custom post-processing pipeline to identify motion and global signal outlier timepoints (global signal>5*σ*, motion derivative >.9mm), regress out tissue-specific nuisance time series (5 eigenvectors each from eroded white matter and CSF masks^56^) and motion-related time series (24 total, including 6 motion parameters, their backward derivatives, and the squares of both^57^), temporally filter (high-pass >0.01 Hz for HCP-YA, >0.008 Hz for HCP-A/D, using DCT projection), and parcellate time series using each atlas. Outlier timepoints were excluded from nuisance regression and temporal filtering. Regional time series for each brain region were obtained by averaging voxels in the denoised time series data. Regional time series for each of the four (for HCP-YA) or two (for HCP-Lifespan) scans were variance-normalized and concatenated, and Pearson correlation between regional time series (excluding outlier timepoints) resulted in an region×region resting-state functional connectivity (FC) matrix for each subject. We also computed an FC matrix with global signal regression, FC_gsr_, by regressing the mean gray matter time series and its temporal derivative from each regional time series before computing region×region Pearson correlation. Finally, we computed a Tikhonov-regularized partial correlation FC_pcorr_ (or FCpc) that minimizes the average distance between subject FC_pcorr_ and the unregularized population average partial correlation^36^; see **Supplemental Methods** for regularization details.

### 4.3 Construction of Structural Connectomes

Diffusion MRI were preprocessed by HCP using the Minimal Preprocessing Pipeline, which jointly corrected for motion, EPI distortion, and eddy current distortion using FSL’s “topup” and “eddy” tools^58, 59^, before being linearly coregistered to the anatomical image. Preprocessed diffusion MRI were further processed using MRtrix3^60^, including bias correction, constrained spherical deconvolution (multi-shell, multi-tissue FOD estimation, lmax=8^61^), and whole-brain tractography. We performed both deterministic (SD_STREAM^62^, or SCdt) and probabilistic, anatomically-constrained tractography (iFOD2+ACT^63, 64^, or SCpr) using dynamic white-matter seeding^65^ for both methods, resulting in 5 million total streamlines per subject for each tractography algorithm. Structural connectivity (SC) matrices were constructed for each atlas by counting the number of streamlines that ended in each pair of gray matter regions, normalized by the total volume of each region pair.

### 4.4 Parcellations

We used three different whole-brain atlases, covering a range in size and construction methodology. The 86-region FreeSurfer atlas (FS86) combines 68 cortical gyri from Desikan-Killiany and 18 subcortical gray matter regions (aparc+aseg^66, 67^). The 268-region Shen atlas (Shen268 or sh268) is an MNI-space cortical and subcortical volumetric atlas based on resting-state fMRI clustering^68, 69^. The 439-region CocoHCP439 atlas (Coco439 or co439) combines 358 cortical regions from the HCP multimodal parcellation^70^, defined using anatomical and functional connectivity gradients, with 12 subcortical regions from FreeSurfer “aseg” (further modified by FSL FIRST^71^), 30 thalamic nuclei derived from FreeSurfer7^72^ (50 original outputs included many small nuclei, which were merged into the final set of 30), 27 cerebellar regions from the SUIT atlas^73^, and 12 additional subcortical nuclei from AAL3^74^. The FS86 and Coco439 atlases were defined based on each individual subject’s FreeSurfer surface and subcortical parcellations, whereas the Shen268 atlas was applied directly to MNI-resampled data for each subject. The FS86, Shen268, and Coco439 parcellations result in 3655, 35778, and 96141 pairwise connectivity estimates, respectively.

### 4.5 Evaluating connectome predictions

As previously discussed, and as illustrated in **Fig. S1**, we consider several complementary assessments of how predicted connectomes (denoted 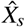 for subject *s*) reflect measured connectomes (denoted *X*_*s*_). Based on these considerations, we use the following metrics for evaluating model performance:

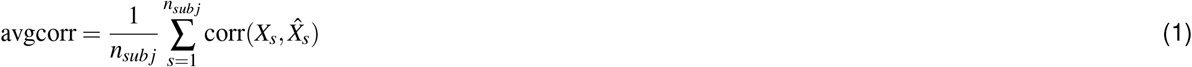

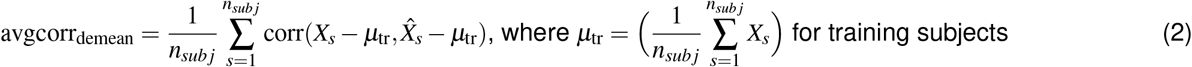

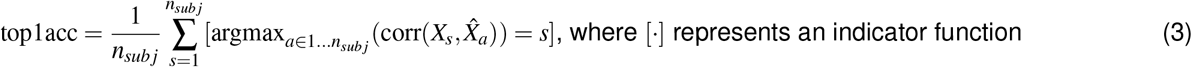

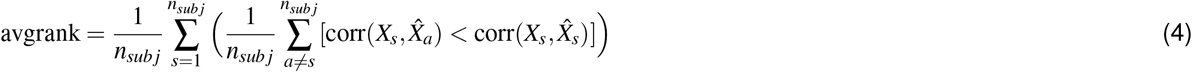

### 4.6 Model training

Each model training epoch comprises two stages: 1) path-wise reconstruction and inter-subject dispersion and 2) latent space consistency. For a given path *i* → *j*, we loop through batches of subjects (*n*_*batch*_ = 41), apply *Encoder*_*i*_ to transform input flavor 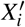 (256 × *n*_*batch*_) to latent representation *z*_*i*_ (128 × *n*_*batch*_), and *Decoder*_*j*_ to transform *z*_*i*_ to predicted output flavor 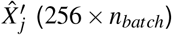. Path-wise reconstruction loss and inter-subject dispersion terms (*L*_*r*.*corrI*_, *L*_*r*.*mse*_, *L*_*r*.*marg*_) are computed per batch from 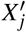 and 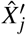. These reconstruction losses collectively penalize reconstruction error in both absolute MSE and correlation terms, and promote inter-subject separation in the reconstruction space. Path-wise latent space dispersion loss (*L*_*z*.*corrI*_, *L*_*z*.*dist*_), computed from *z*_*i*_, promotes intersubject separation in the latent space. This forward pass and backpropagation is repeated for each batch of training subjects, before moving on to the next training path. The order in which paths are selected is randomly reshuffled each epoch. The second stage of each training epoch, latent space consistency, enforces similarity of the latent space representations from each input flavor. We re-compute *z*_*i*_ for all input flavors from the encoders updated in the path-wise stage, and then compute *L*_*z*.*sim*_ as the total MSE between the set of *z*_*i*_ matrices for all flavors.

Explicit weights were applied to the reconstruction MSE loss term *L*_*r*.*mse*_ (*w*_*rm*_ = 1000), the combined path-wise latent space dispersion loss *L*_*z*_ (*w*_*z*_ = 10), and the latent space intra-subject consistency loss *L*_*z*.*sim*_ (*w*_*zs*_ = 10000). See **Algorithm 1** for pseudocode depiction of training procedure, and **Table 1** for details of loss function terms.

After each training epoch, we compute predicted connectomes for all paths on a held-out set of 80 validation subjects, and the inverse PCA transform (pre-computed on the training data only) is applied to compute 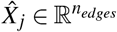 from 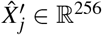. Reconstruction accuracy and identifiability are computed in this high-dimensional data space. Loss term weights were tuned empirically to balance the reconstruction (avgcorr_demean_) and identifiability (avgrank) performance metrics in the validation dataset. **Fig. S2** illustrates this trade-off when tuning MSE reconstruction loss.

The model was implemented in PyTorch 1.10, using the AdamW optimizer with learning rate 10^−4^. Model weights were regularized by weight decay 10^−2^, as well as dropout in all encoders and decoders (dropout rate 0.5). The model was trained for 5000 epochs, which took approximately 36 hours on an Nvidia A100 GPU. Due to the regularization and the largely linear nature of the model, we did not observe significant overfitting. Validation performance plateaued after 500-1000 epochs, and we selected the 2000 epoch checkpoint for further evaluation.

#### Algorithm 1 Krakencoder model training procedure. (See Table 1 for loss function details)

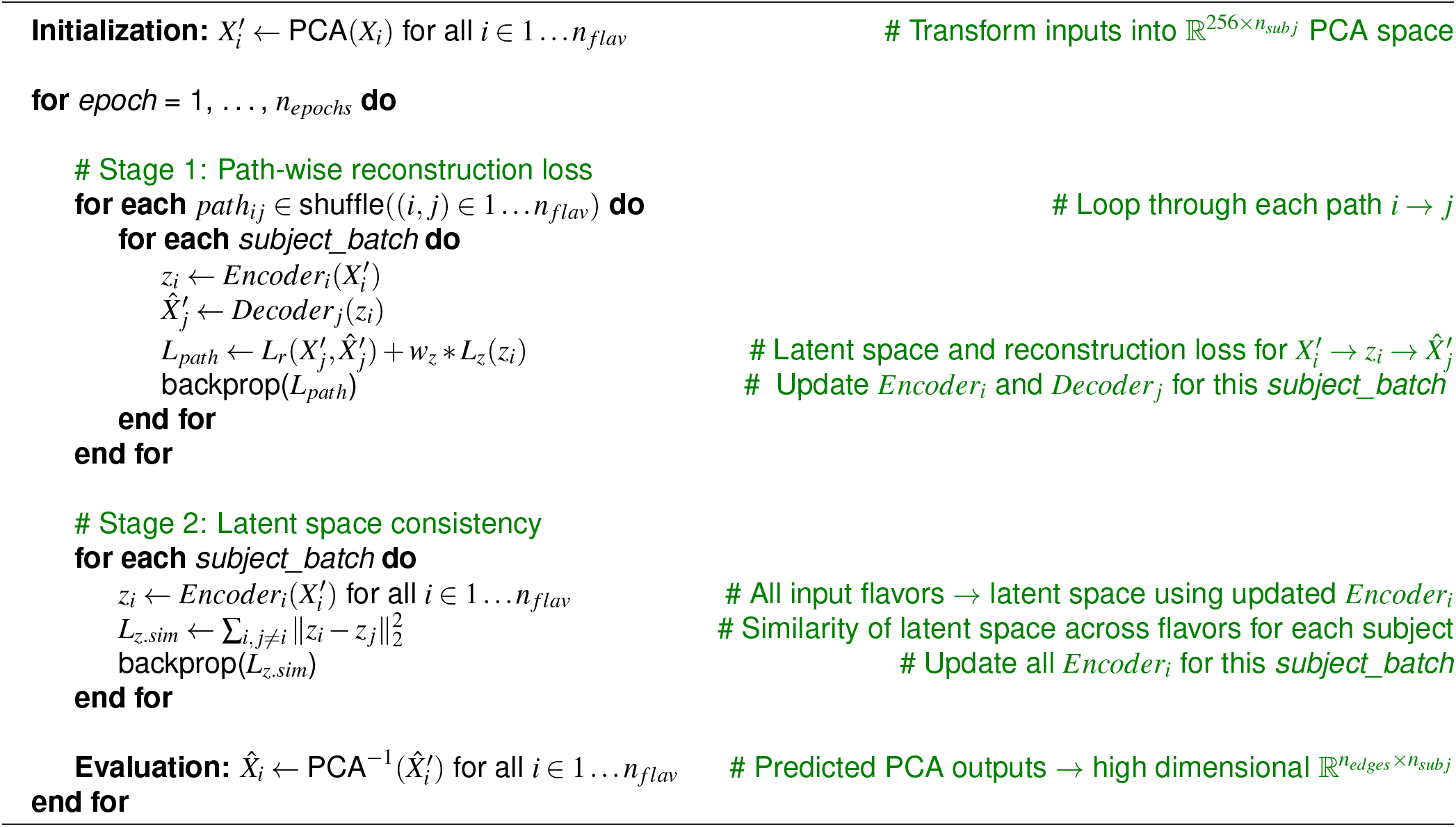

### 4.7 Procedure for comparing family structure and predicting demographics

To examine how the connectomes (predicted and measured) reflect family structure, we first compute the averaged latent vector in the Krakencoder for each subject from either all 15 flavors (“fusion”) or the 9 FC (“fusionFC”) and 6 SC (“fusionSC”) separately, and then compute a *n*_*sub j*_ × *n*_*sub j*_ inter-subject latent similarity matrix using Pearson correlation. These pairwise similarities are grouped according to the inter-subject relationship. For comparison, we also compute the inter-subject similarity for all 15 flavors of the high-dimensional measured connectome data, average these 9, 6, or 15 *n*_*sub j*_ × *n*_*sub j*_ similarity matrices, and group them according to relatedness. Due to the high baseline similarity in observed data, latent and observed data similarity measures were independently z-scored by the mean and standard deviation of the similarity of unrelated subjects for visual comparison. Differences of ROC separability of the distributions (e.g., separability of DZ and MZ distributions in latent space, compared to separability of DZ and MZ in observed data) was assessed through permutation testing (10000 matched permutations).

For demographic predictions, we used a linear support vector classifier (SVC) to predict sex, and kernel ridge regression with linear kernel to predict age and cognition^75^. For the Krakencoder, models were fit using crossvalidation on the average latent vectors for the 683 subjects used to train the Krakencoder model, and tested on the 196 held-out subjects. For the observed connectome data, we computed the inter-subject cosine similarity matrix of each connectome flavor, and used the averaged similarity matrices as input into the model, using the same training and testing splits as for the Krakencoder. Regression hyperparameters were selected by nested cross-validation grid search. Variability of prediction accuracy was assessed through bootstrap resampling (N=100), and significant differences between Krakencoder and observed data models was assessed using permutation testing of matched bootstrap samples (10000 permutations).

## Data and code availability

Code is available at https://github.com/kjamison/krakencoder. Post-processing code for FC can be found here: https://github.com/kjamison/fmriclean. Additional data files are available at https://osf.io/dfp92.

Preprocessed data for this study are available for download from the Human Connectome Project (www.humanconnectome.org). Users must agree to data use terms for the HCP before being allowed access to the data and ConnectomeDB, details are provided at https://www.humanconnectome.org/study/hcp-young-adult/data-use-terms. Data from the HCP Aging and HCP Development studies can be downloaded as part of the HCP-Lifespan 2.0 release, distributed by the NIMH Data Archive (https://nda.nih.gov). See https://www.humanconnectome.org/study/hcp-lifespan-aging/data-releases and https://www.humanconnectome.org/study/hcp-lifespan-development/data-releases for more information about data use terms.

Post-processed connectomes and related input files can be made available upon reasonable request from corresponding author K.J., subject to HCP data use restrictions described above.

## Acknowledgements

This work was supported by the following grants: NIH RF1 MH123232 (AK), Ann S. Bowers Foundation (AK), NIH R01 AG053949 (MS), NSF CAREER 1748377 (MS).

Data were provided by the Human Connectome Project, WU-Minn Consortium (Principal Investigators: David Van Essen and Kamil Ugurbil; 1U54MH091657) funded by the 16 NIH Institutes and Centers that support the NIH Blueprint for Neuroscience Research; and by the McDonnell Center for Systems Neuroscience at Washington University.

Research reported in this publication was supported by the National Institute Of Mental Health of the National Institutes of Health under Award Number U01MH109589, National Institute On Aging of the National Institutes of Health under Award Number U01AG052564. The HCP-Development 2.0 Release data used in this report came from DOI: 10.15154/1520708. The HCP-Aging 2.0 Release data used in this report came from DOI: 10.15154/1520707.

The authors would like to thank Fang-Chang Yeh for sharing pre-processed diffusion MRI data for HCP-Development and HCP-Aging subjects.

## Author contributions statement

K.W.J. conceived the study and developed the software with help from Z.G.. K.W.J., A.K., and M.R.S. analyzed results, and Q.W. and C.T. helped with validation. K.W.J., A.K., and M.R.S. wrote and edited the manuscript. All authors reviewed the manuscript.

## Competing interests

The authors declare no competing interests.

## Notes

### Competing Interest Statement

The authors have declared no competing interest.

### Summary of Updates

New sensitivity analysis, adding new connectome types, and validating on clinical data

