## Supplemental Information for "Release the Krakencoder: A unified brain connectome translation and fusion tool"

### Supplemental Information for
### *Release the Krakencoder: A unified connectome translation and fusion tool*

#### Supplemental Methods

| Dataset | N | Demographics | | Acquisition |
|---|---|---|---|---|
| HCP-YA (training and testing) | 958 | Age 22-37<br>442 M (27.9±3.7)<br>516 F (29.5±3.6)<br><br>40 retest sessions<br>11 M (27.3±3.8)<br>29 F (31.8±1.9) | 115 MZ twin pairs<br>43×2 M (27.8±3.5)<br>72×2 F (30.4±2.8)<br><br>64 DZ twin pairs<br>29×2 M (26.9±3.5)<br>35×2 F (30.7±3.1) | **Resting-state fMRI**: 2mm vox, TR 720ms, LR/RL phase-encoding, **1 hour total**<br>**Diffusion MRI**: 1.25mm vox, b=1000,2000,3000, 90dir/shell, LR/RL phase-encoding, **1 hour total** |
| HCP-D (testing) | 608 | Age 8-21<br>282 M (14.8±3.8)<br>326 F (14.6±4.0) | 76 sibling pairs (50 MM/FF, 26MF, Δage 0-9yr) | **Resting-state fMRI**: 2mm vox, TR 800ms, AP/AP phase-encoding, **25.5 min total**<br>**Diffusion MRI**: 1.5mm vox, b=1500,3000, 90dir/shell, AP/PA phase-encoding, **22.5 min total** |
| HCP-A (testing) | 716 | Age 36-100+<br>312 M (60.8±15.6)<br>404 F (60.1±15.7 | | |
| Multiple Sclerosis (testing) | 100 | Age 22-71<br>34 M (47.4±10.8)<br>66 F (44.8±12.1) | Patients with white-matter lesions | **Resting-state fMRI**: 3.75x3.75x4mm vox, TR 2310ms, AP phase-encoding, **6.9 min total**<br>**Diffusion MRI**: 1.8x1.8x2.5mm vox, b=800, 64dir, AP phase-encoding, **9.75 min total** |

**Table S1.** Summary of datasets used in this study

***Regularized partial correlation for functional connectivity***

Following Pervaiz[1], we identify a regularization parameter $\lambda$ that minimizes the euclidean norm between the regularized precision matrices and the population average of the unregularized precision matrices:

$$\overline{\Omega} = \frac{1}{n_{subj}} \sum_{a}^{n_{subj}} FC_a^{-1}$$
1. Compute population mean unregularized inverted $FC$

$$\hat{\Omega}_a = [FC_a + \lambda I]^{-1}$$
$$\lambda = \text{argmin}_\lambda \sum_{a}^{n_{subj}} \|\hat{\Omega}_a - \overline{\Omega}\|_2$$
2. Find $\lambda$ that minimizes MSE loss between subject $\hat{\Omega}$ and target $\overline{\Omega}$

$$FC_{pcorr}(i,j) = -\hat{\Omega}_{ij} / \sqrt{\hat{\Omega}_{ii}\hat{\Omega}_{jj}}$$
3. Normalize precision matrix $\hat{\Omega}$ to compute partial correlation $FC_{pcorr}$

For Shen268 and Coco439 atlases, the target $\overline{\Omega}$ was computed by averaging the pseudoinverse of each subject FC rather than inverse. A different $\lambda$ was identified for each atlas, based on the 700 training subjects from HCP-YA. Optimal $\lambda$ was 0.06 for FS86, 0.15 for Shen268, and 0.25 for Coco439. For HCP-Aging and HCP-Development, regularization target $\overline{\Omega}$ was the $\overline{\Omega}$ from HCP-YA training data. For HCP-Aging: FS86 $\lambda$=0.10, Shen268 $\lambda$=0.31, Coco439 $\lambda$=0.57. For HCP-Development: FS86 $\lambda$=0.17, Shen268 $\lambda$=0.35, Coco439 $\lambda$=0.54.

#### Supplemental Figures

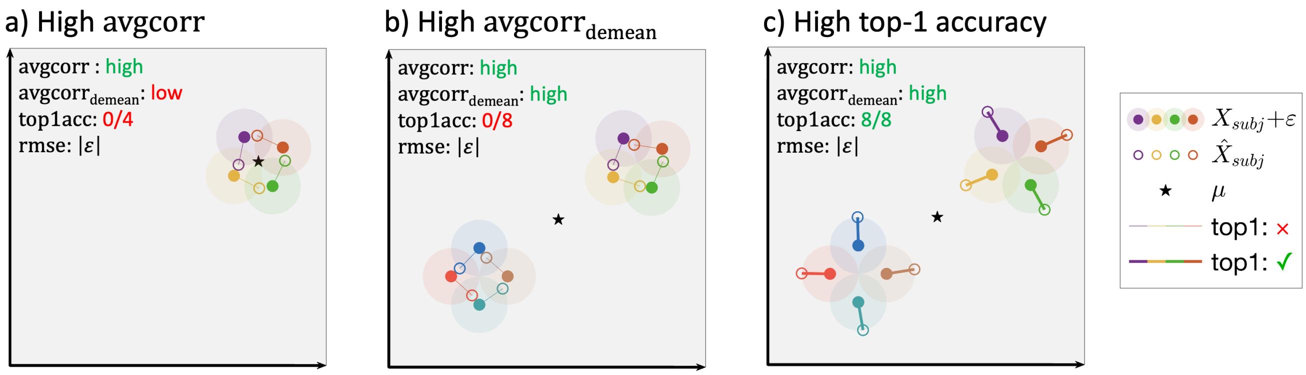


**Figure S1. Connectome prediction similarity and identifiability can be complementary.** Filled circles represent each subject's measured connectome ($X_{subj}$), open circles represent each subject's predicted connectome ($\hat{X}_{subj}$), and the black star is the population mean of all $X_{subj}$. Lines connecting each subject's measured and predicted connectomes are thick when that subject's $X_{s=a}$ is closer to its own $\hat{X}_{s=a}$ than any other $\hat{X}_{s\neq a}$. All three visualized predictions have the same residual magnitude $|\varepsilon|$ (i.e., RMSE), and similar average predicted correlation, but have very different measured→predicted identifiability: Predictions for a given subject ($\hat{X}_{subj}$) can be closer to the measured data of another subject than their own measurement ($X_{subj}$). **a.** Average correlation between $X_{subj}$ and prediction $\hat{X}_{subj}$ (avgcorr) is high because inter-subject similarity is high, but every subject's prediction is closer to another subject's measurement than their own. **b.** Removing the population mean $\mu$ before computing correlation ($\mathrm{avgcorr}_{\mathrm{demean}}$) does not resolve this conflict when data are not uniformly distributed, for example when clustered. **c.** Predictions with the same RMSE and similar correlation as in the other panels, but with perfect top-1 identifiability. Thus, accuracy metrics such as MSE or correlation do not guarantee a precise match, which should instead be measured explicitly through Top-1 accuracy or average rank percentile.

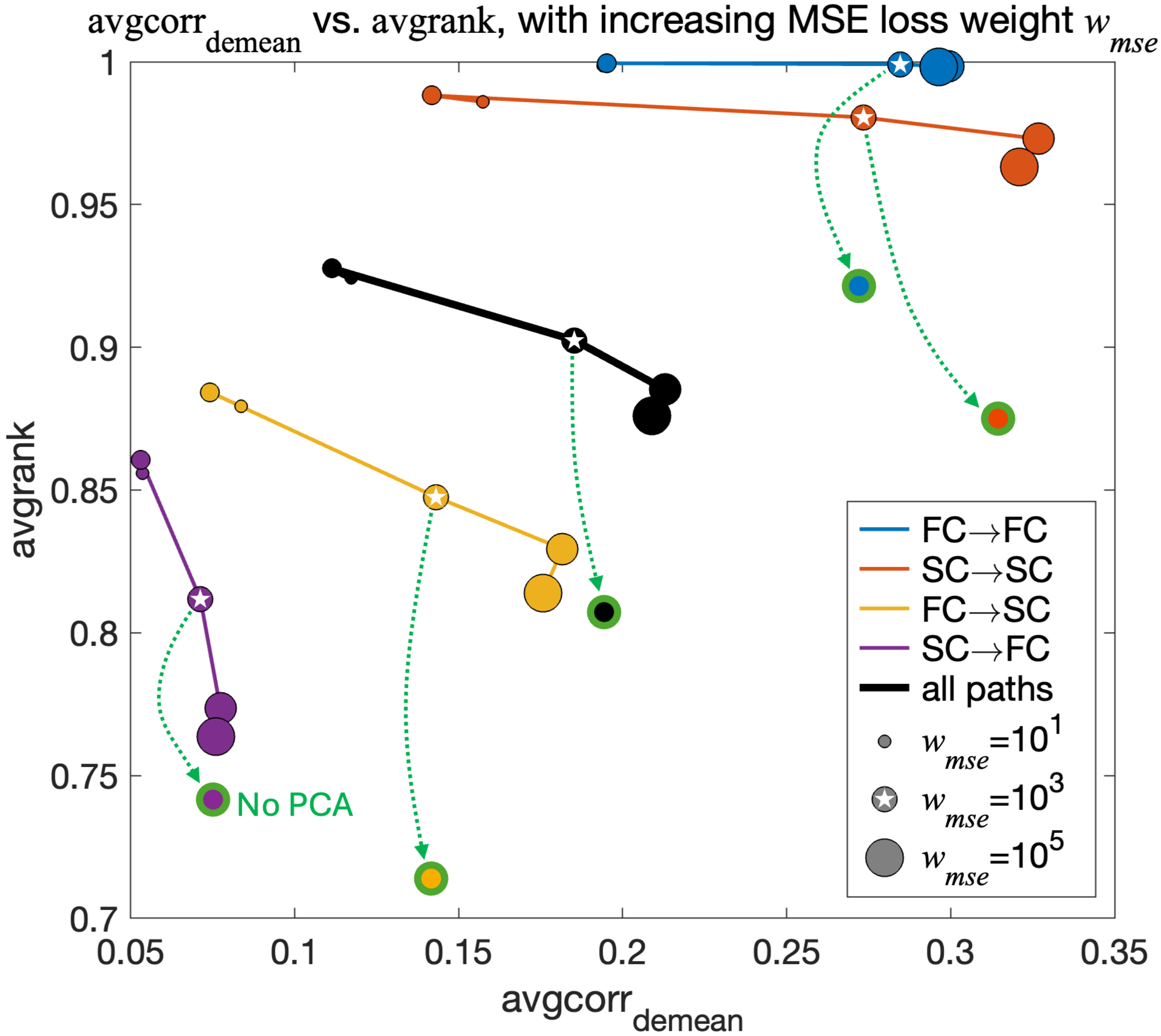


**Figure S2. Increasing reconstruction loss weight improves reconstruction accuracy but decreases identifiability.** Model loss weights were tuned to optimize the reconstruction accuracy, measured by $\mathrm{avgcorr}_{\mathrm{demean}}$, and reconstruction identifiability, measured by $\mathrm{avgrank}$, on validation subjects. Each point shows the performance averaged across prediction categories for a model trained with a different reconstruction loss weight $w_{mse} = 1 - 100000$. The intra-modality predictions (SC→SC in red and FC→FC in blue) have consistently high identifiability ($\mathrm{avgrank}$), and increasing reconstruction loss weight $w_{mse}$ increases reconstruction accuracy ($\mathrm{avgcorr}_{\mathrm{demean}}$) without decreasing identifiability. Inter-modality predictions (FC→SC in yellow and SC→FC in purple), have lower accuracy and identifiability, and SC→FC identifiability decreases sharply with increasing $w_{mse}$. Reconstruction loss weight $w_{mse} = 1000$ was chosen to balance these two factors. Performance of a model trained without using PCA dimensionality reduction (green outlines) shows reduced identifiability without improving $\mathrm{avgcorr}_{\mathrm{demean}}$.

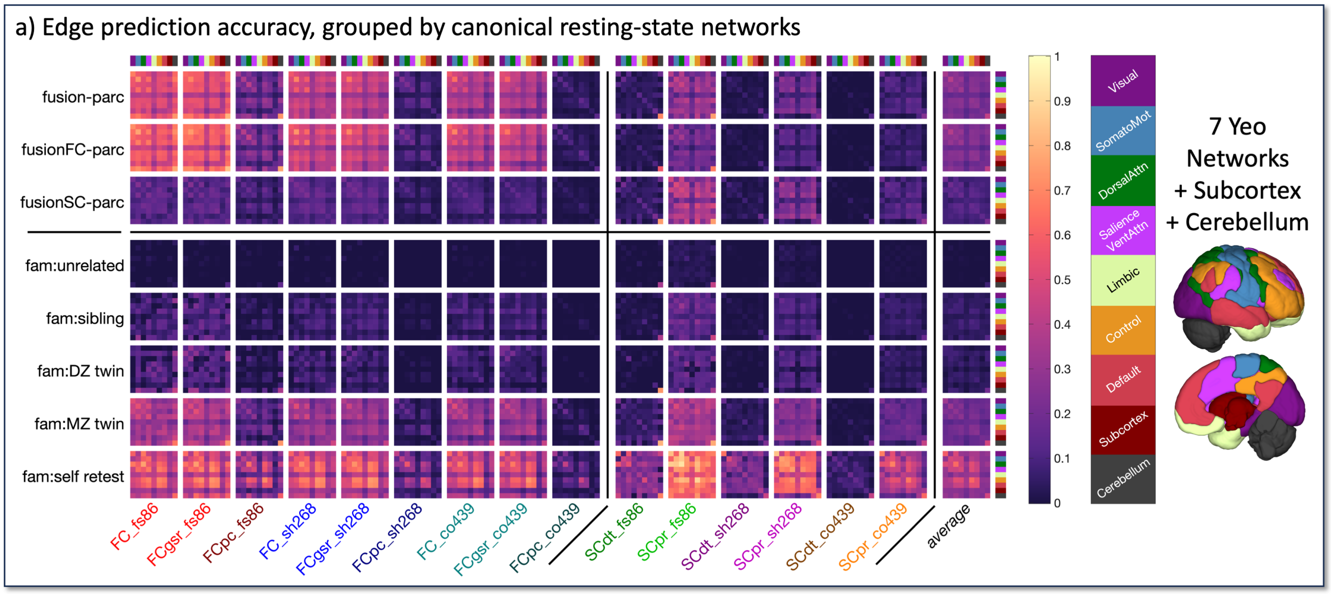


b) Similarity of network-grouped spatial pattern to self-retest edge reliability baseline

| | FC_fs86 | FCgsr_fs86 | FCpc_fs86 | FC_sh268 | FCgsr_sh268 | FCpc_sh268 | FC_co439 | FCgsr_co439 | FCpc_co439 | SCdt_fs86 | SCpr_fs86 | SCdt_sh268 | SCpr_sh268 | SCdt_co439 | SCpr_co439 | average |
|---|---|---|---|---|---|---|---|---|---|---|---|---|---|---|---|---|
| fusion-parc | .48 | .58 | .80 | .62 | .68 | .95 | .76 | .79 | .89 | .70 | .61 | .77 | .89 | .86 | .81 | .75 |
| fusionFC-parc | .50 | .60 | .79 | .61 | .65 | .94 | .76 | .78 | .89 | .66 | .53 | .77 | .90 | .88 | .84 | .74 |
| fusionSC-parc | .50 | .57 | .95 | .83 | .90 | .98 | .81 | .83 | .91 | .73 | .64 | .77 | .88 | .85 | .77 | .79 |
| fam:unrelated | .00 | -.01 | .22 | .08 | -.02 | .18 | -.22 | -.26 | -.03 | .51 | .01 | .47 | .65 | .53 | .69 | .19 |
| fam:sibling | .21 | .29 | .42 | .58 | .56 | .87 | .66 | .67 | .92 | .62 | .53 | .70 | .88 | .80 | .85 | .64 |
| fam:DZ twin | .11 | .11 | .51 | .51 | .52 | .86 | .64 | .54 | .90 | .52 | .36 | .57 | .82 | .69 | .75 | .56 |
| fam:MZ twin | .47 | .61 | .71 | .64 | .79 | .97 | .83 | .90 | .96 | .67 | .43 | .77 | .84 | .86 | .83 | .75 |
| fam:self retest | 1 | 1 | 1 | 1 | 1 | 1 | 1 | 1 | 1 | 1 | 1 | 1 | 1 | 1 | 1 | 1 |

**Figure S3. Spatial variation in edge prediction accuracy and baseline reliability.** For the Krakencoder's fusion predictions made without the parcellation being predicted (fusion-parc, fusionFC-parc, and fusionSC-parc), the prediction accuracy for each edge is the Pearson correlation between the predicted and measured values for 196 held-out test subjects. To simplify visualization and compare across parcellations, each region is assigned to one of 7 cortical resting state networks[2] or separate subcortical and cerebellar networks, and the edge-wise accuracies (region $\times$ region) are then grouped and averaged into a $9 \times 9$ network representation. Age- and sex-matched individuals of given relatedness, as well as self-retest, are correlated and averaged in the same manner, in order to provide a baseline for comparing the spatial patterns of edge-wise accuracy. **a.** The spatial heatmaps show that prediction accuracy is relatively uniform throughout the cortex and subcortex. Networks with lower predictability (e.g., the limbic network), also show low reliability in self-retest measurement reliability (bottom row). **b.** Each network $\times$ network heatmap in **a** is correlated with the corresponding self-retest (bottom row), to show the consistency of the spatial pattern of edge-wise predictions.

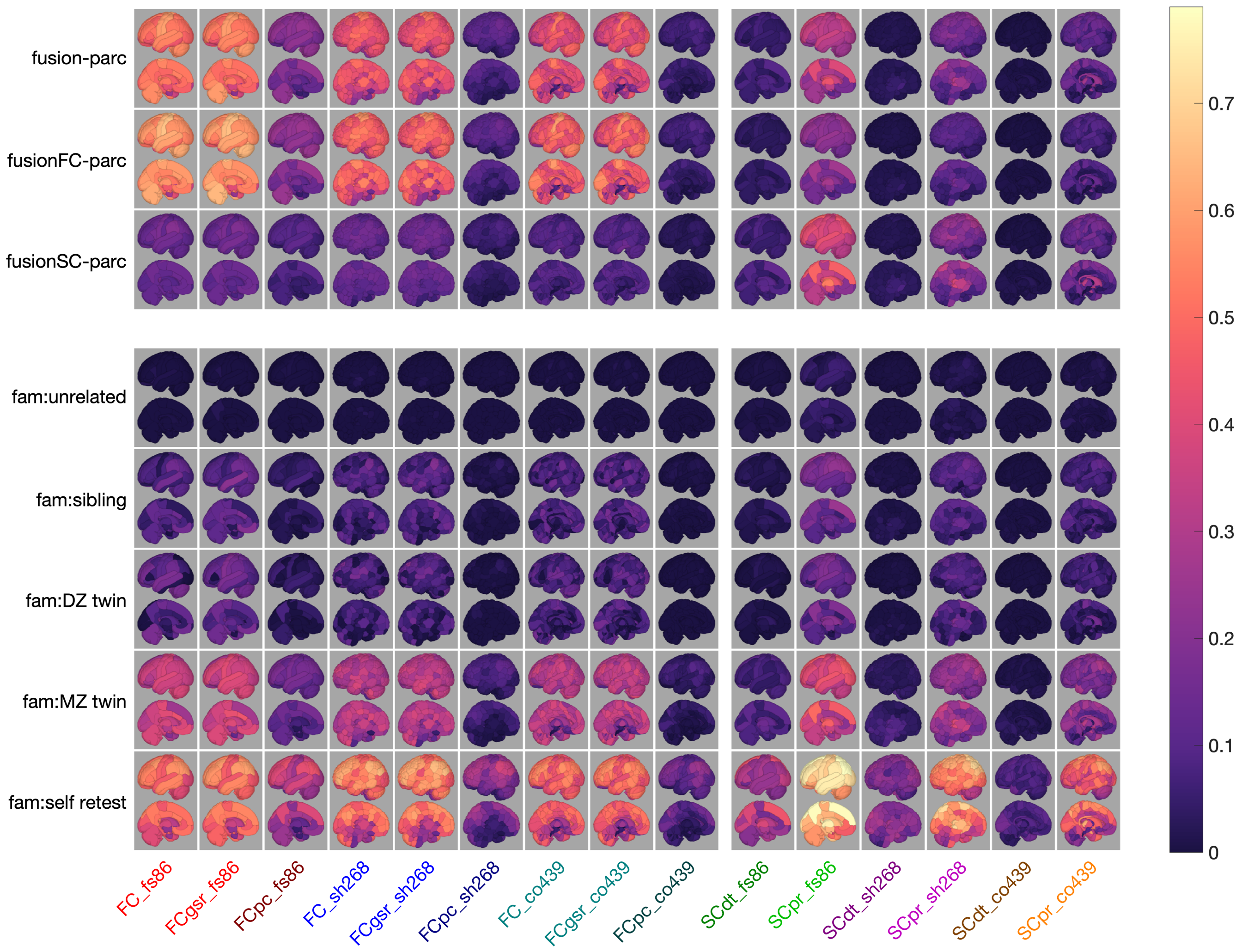


**Figure S4. Regional variation in prediction accuracy and baseline reliability.** For the Krakencoder's fusion predictions made without the parcellation being predicted (fusion-parc, fusionFC-parc, and fusionSC-parc), the prediction accuracy for each region is the Pearson correlation between the predicted and measured edge values for 196 held-out test subjects, averaged across all other regions. Age- and sex-matched individuals of given relatedness, as well as self-retest, are correlated and averaged in the same manner, in order to provide a baseline for comparing the spatial patterns of edge-wise accuracy. Only the left hemisphere is shown, to simplify visualization.

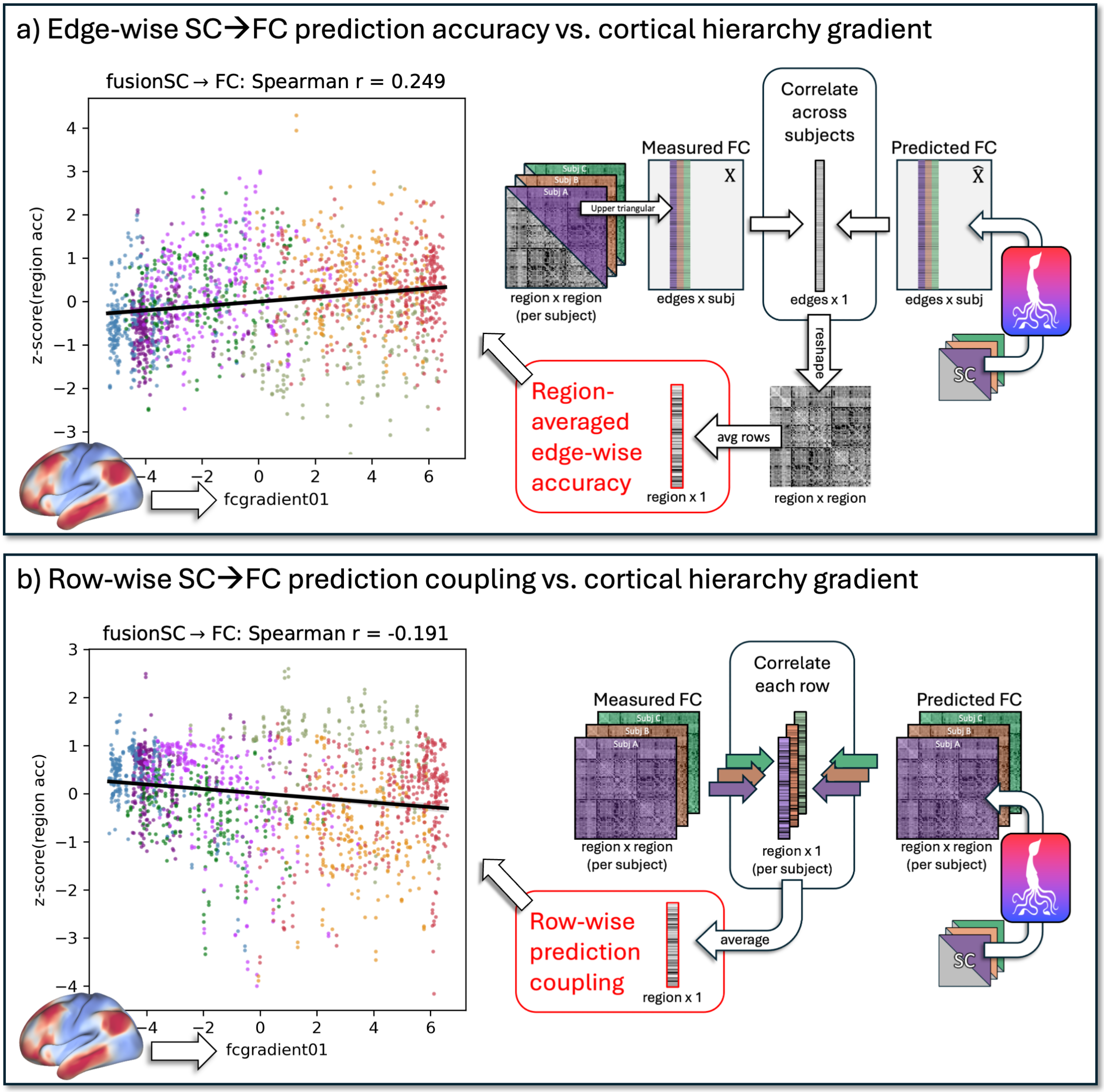


**Figure S5. Inter-modality prediction accuracy aligns with cortical hierarchy.** Regional SC→FC prediction accuracy covaries with the principle gradient of cortical hierarchy derived from functional connectivity[3]. **a.** Edge-wise prediction accuracy, measuring the explained inter-subject variance for each connectome edge, is highest for high-level association cortex (red areas on the cortical map, including regions in default mode and control networks). **b.** Conversely, estimates of regional coupling, computed by correlating each region's row in the measured and predicted connectome, are highest for low-level sensory cortex (blue areas on the cortical map, including regions in visual and somatomotor networks). Each dot in the scatter plot represents a region colored according to its assigned Yeo network[2].

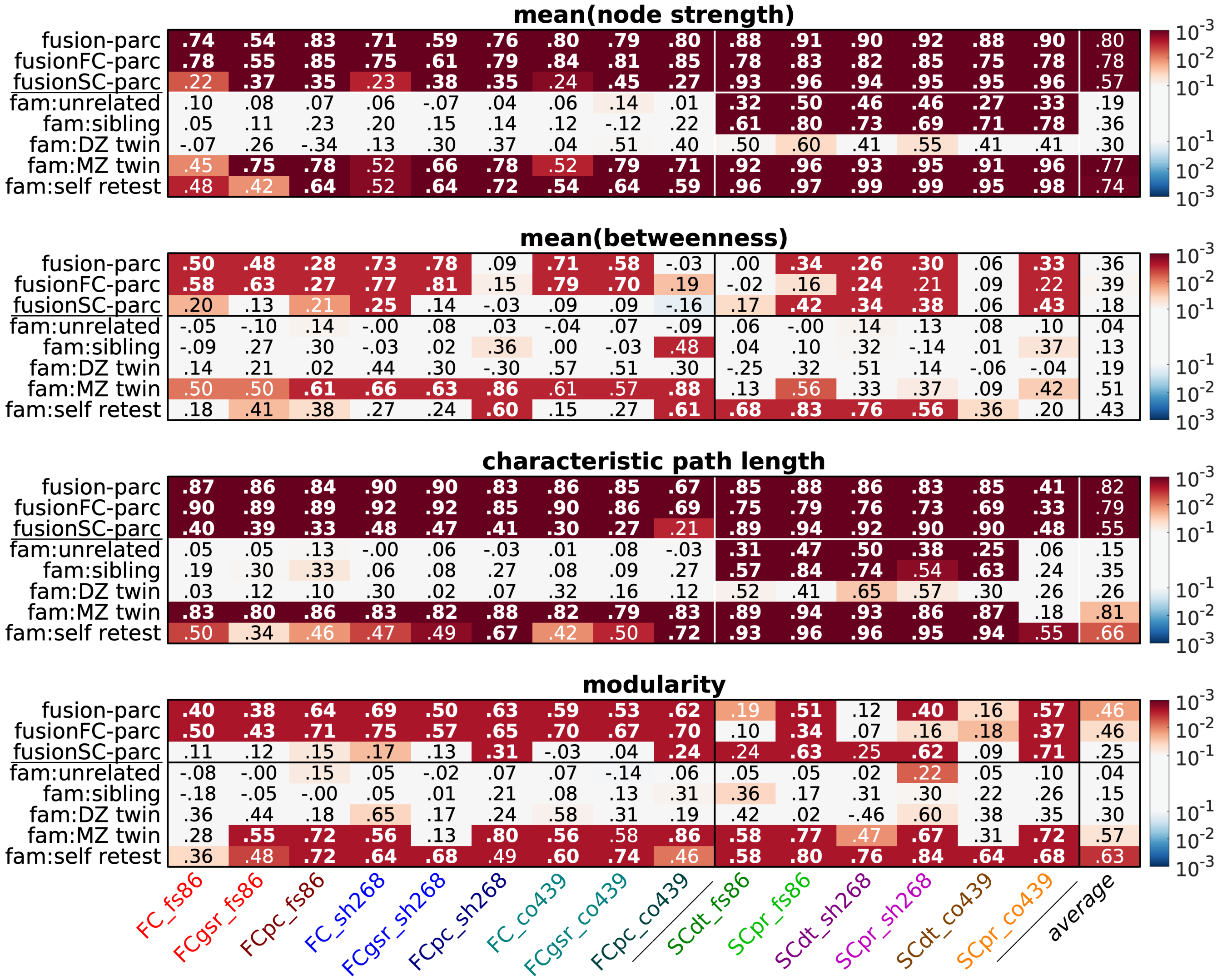


**Figure S6. The Krakencoder preserves network properties of the connectomes.** Each numbered entry in the heatmap shows the Spearman rank correlation between the graph metric of the Krakencoder's predicted connectome (based on one of the three fusion representations, "fusion-parc", "fusionFC-parc" and "fusionSC-parc", where inputs exclude the parcellation being predicted) for that output connectome flavor listed on the x-axis, compared to the graph metric for that measured flavor of connectome. These values were calculated using the held-out set of 196 individuals. Colors indicate the correlation's p-value. Bold entries denote significant correlation ($p_{perm} < 10^{-3}$, 1000 permutations, FDR-corrected).

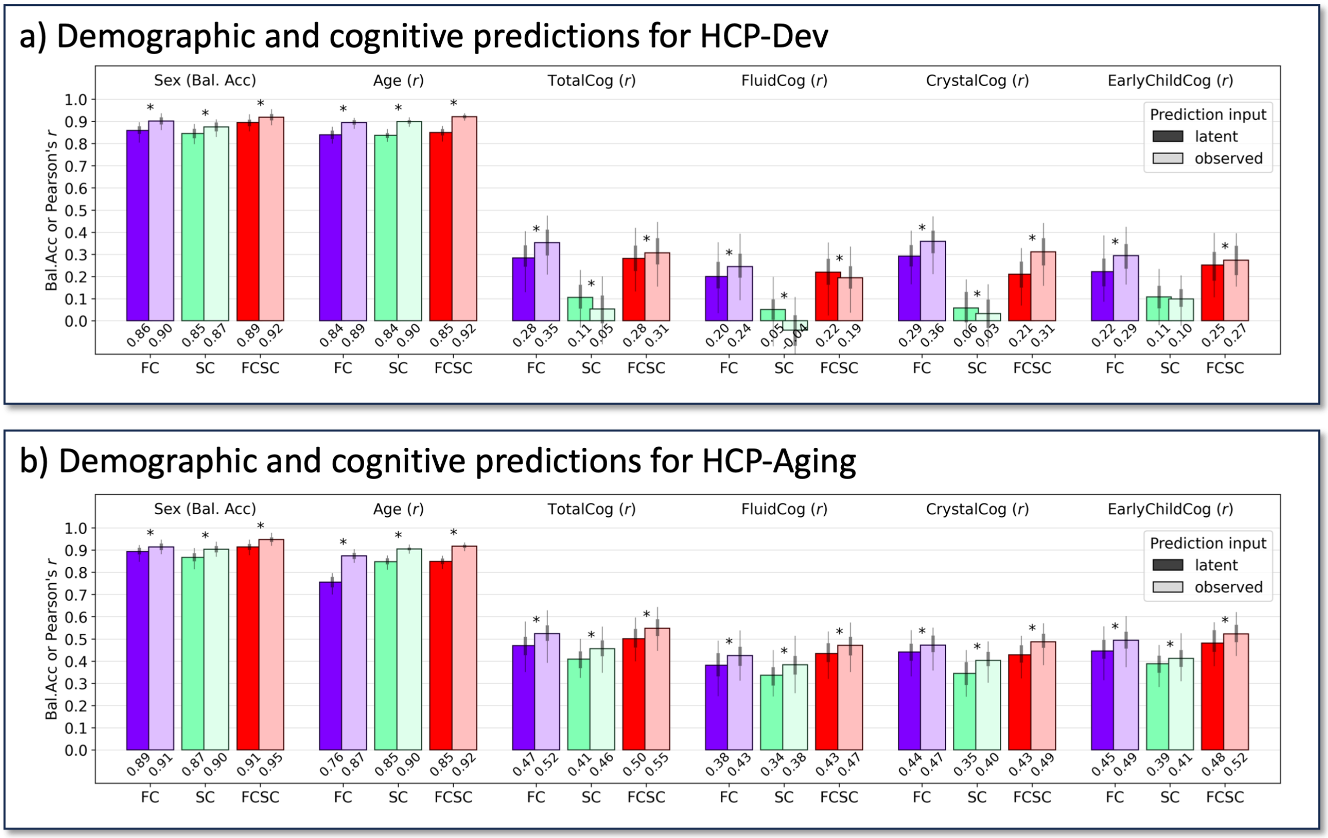


**Figure S7. Demographic prediction for HCP-Lifespan (out-of-sample test set).** Prediction accuracy of demographic and cognitive metrics from different subsets of the latent space and observed connectome data, for HCP-Dev (**a**) and HCP-Aging (**b**). Accuracy of predictions using observed data exceeds that from latent space. See **Fig. 3b** for more information on predictions. $*=$ significant difference between prediction accuracy from latent space and observed data. ($p_{perm} < 10^{-3}$, 1000 permutations, FDR-corrected).

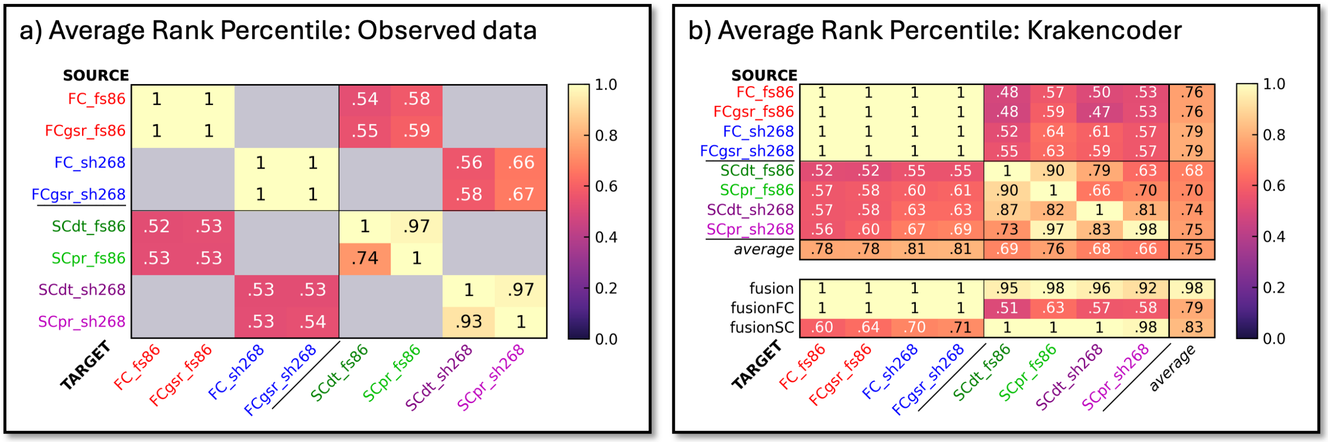


**Figure S8. Krakencoder performance in individuals with multiple sclerosis (out-of-sample and out-of-distribution test set).** Average rank percentiles of identifiability in 100 individuals with multiple sclerosis using (**a**) the observed SC and FC (within atlas comparisons only) and (**b**) the Krakencoder's individual arm predictions (top) and the fusion/fusionSC/fusionFC predictions (bottom). Note that the identifiability of the SC to FC mapping in the observed data is just above chance levels ($\leq$ 54%) but the Krakencoder's fusionSC to FC predictions are as high as 71%.

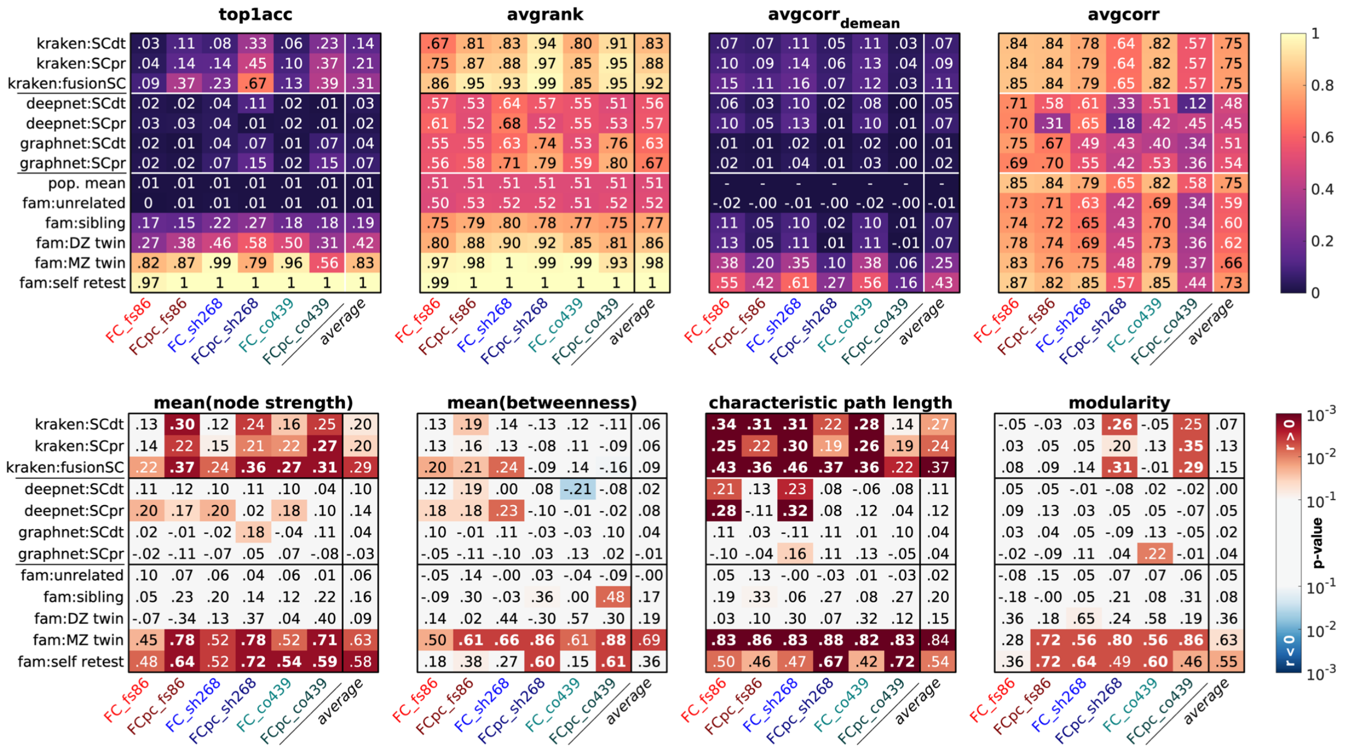


**Figure S9. Comparison of the Krakencoder's SC to FC mapping to state-of-the-art ML-based SC to FC mapping.** Here we compare the Krakencoder's performance in mapping from SC to FC, which is the most common mapping in previous work. We compare against a deep neural network ("deepnet")[4] and a deep graph neural network ("graphnet")[5]. Top-1 accuracy, average rank percentile, average correlation of predicted and measured FC after and before de-meaning are presented in the top row of panels, one value for each FC flavor as output (columns) and various SCs as input (rows). Network metrics are provided in the bottom row of figure panels. There are three SC inputs for the Krakencoder model: volume-normalized deterministic SC for that parcellation only [kraken:SCdt], volume-normalized probabilistic SC for that parcellation only [kraken:SCpr] and the fusion of all 6 SC inputs in the latent space [kraken:fusionSC]. There are two SC input flavors (deterministic [SCdt] and probabilistic [SCpr] volume normalized SC) for the deepnet and graphnet models. The same metrics between pairs of measured FC from varied familial relatedness categories is shown as a comparison. Deepnet and graphnet models were trained and evaluated using the same data and subject splits as the Krakencoder model.

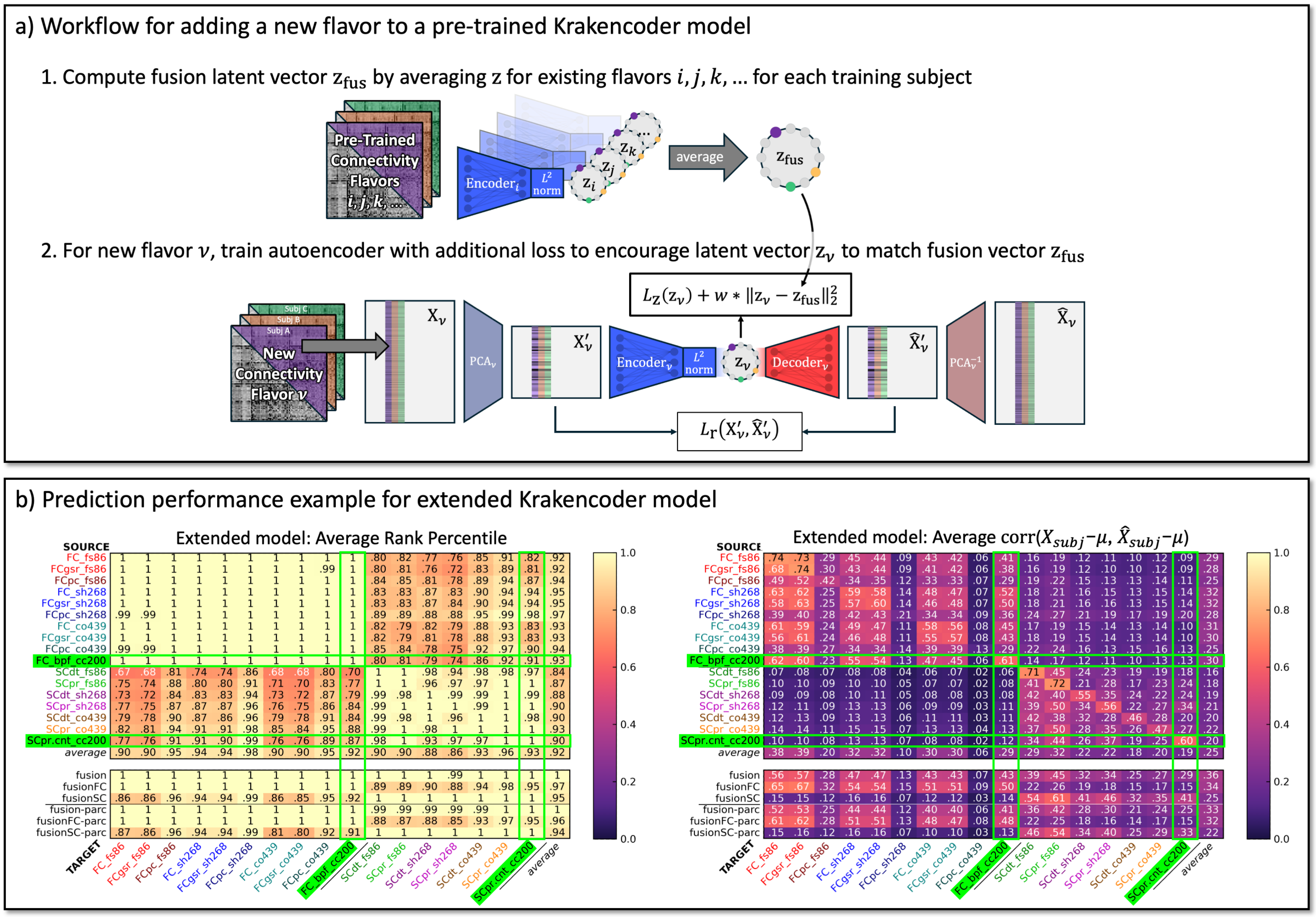


**Figure S10. A pre-trained Krakencoder model can be extended to new flavors. a.** To add a new flavor, we first compute the fusion representation $z_{\text{fus}} \in \mathbb{R}^{128 \times n_{subj}}$ by averaging the latent vector for each flavor in the existing model. With new connectome data $X_v \in \mathbb{R}^{n_{edges} \times n_{subj}}$, we compute new PCA transformations and train a new autoencoder, using the same reconstruction and latent loss terms $L_r$ and $L_z$ as the original training, but with an additional loss term to force the new latent $z_v$ to be similar to $z_{\text{fus}}$. **b.** Connectome prediction performance after adding two new flavors to the original 15 flavor model. New flavors shown here use a previously unseen parcellation (cc200[6]), bandpass-filtered time series for FC (original flavors use a high-pass filter), and un-normalized streamline count for SC (original flavors use pairwise volume normalization). Prediction identifiability ($\text{avgrank}$, left) and reconstruction accuracy ($\text{avgcorr}_{\text{demean}}$, right) are comparable to the original flavors for these new flavors, highlighted in green.
